# Biomimetic biaxial mechanical properties enhance hemodynamic performance and prevent adverse tissue remodelling in tissue-engineered heart valves

**DOI:** 10.1101/2025.03.02.641050

**Authors:** Bahram Mirani, Sean O. Matthew, Min Namgung, Neda Latifi, Cassandra Chiao-Ying Yao, Brian G. Amsden, Craig A. Simmons

**Author notes:** Corresponding Authors: Craig A. Simmons Translational Biology & Engineering Program, Ted Rogers Centre for Heart Research University of Toronto 661 University Avenue, 14th Floor Toronto, Ontario Canada, M5G 1M1, Bahram Mirani Translational Biology & Engineering Program, Ted Rogers Centre for Heart Research University of Toronto 661 University Avenue, 14th Floor Toronto, Ontario Canada, M5G 1M1.

## Abstract

Heart valve tissue engineering holds the promise of providing an unlimited supply of heart valve replacements for patients with valvular heart disease. While tissue-engineered heart valves show promising performance in the short term, regurgitation due to fibrotic leaflet shortening remains a persistent limitation in most cases. Here we tested the hypothesis that engineering heart valves with native biaxial mechanical properties would mitigate adverse tissue remodeling in vitro. Melt electrowriting and hydrogel casting were used to generate four types of tissue-engineered heart valves with combinatorial biomimetic or non-biomimetic mechanical properties in the radial and circumferential direction of the heart valves. The heart valves were subject to pulmonary hemodynamic conditions in a pulse duplicator bioreactor, and their tissue remodelling, including cell phenotype and orientation and extracellular matrix properties, was investigated. The results showed that tissue-engineered heart valves with native mechanical properties, particularly in the radial direction, outperformed those with non-physiological mechanics in hemodynamic function, maintaining quiescent cell phenotype and cell orientation, and avoiding fibrosis. This study, for the first time, elucidates the impact of native mechanical properties on the performance of tissue-engineered heart valves and their cell and extracellular matrix homeostasis.

## 1- Introduction

The majority of current tissue-engineered heart valves (TEHVs) fail in the mid-term *in vivo* mainly due to leaflet retraction that leads to valvular insufficiency^1,2^. Despite considerable efforts for almost three decades, this prevalent mode of failure remains largely unsolved and has prevented TEHVs from clinical translation^1,2^. A range of approaches, such as decellularizing valves before implantation^3–9^, modifications to the valve geometry^10,11^, and the use of bioresorbable biomaterials instead of tissues grown in vitro^12–17^, have been taken to mitigate adverse tissue remodelling. Adverse tissue remodelling in TEHVs has also motivated fundamental research on its underlying mechanisms at the cellular and extracellular matrix (ECM) levels. Such studies have primarily relied on computational modelling, where heart valve physiological geometry, mechanics, and boundary conditions along with hemodynamic loading are taken into account^18–20^. Knowing the importance of valve tissue mechanics and boundary conditions in defining the tissue remodelling response (at least through the computational models), experimental studies can better unveil how heart valve tissue cells and ECM components respond to hemodynamic pressures and remodel one another in a reciprocal manner.

Hemodynamic pressure causes a strain response in heart valve tissue that depends on its mechanical properties, geometry, and boundary conditions. This strain response largely regulates cellular behaviour which is responsible for ECM homeostasis^21^. Given that the geometry and boundary conditions of native heart valve tissue are replicated with good accuracy in TEHVs, their deviation from native tissue mechanical properties can be crucial in triggering adverse tissue remodelling. It has been shown that large strain anisotropy in heart valve tissue caused by large circumferential strain and negligible or negative radial strain leads to contractile cell activation^22^ and cell reorientation toward the circumferential direction^18^. This initial tissue remodelling leads to reduced belly curvature in the leaflet tissue, which in turn further reduces the translation of hemodynamic pressure to radial stretch in diastole, resulting in decreased radial strain at larger extents^18^. This results in a positive feedback loop that continues toward heart valve failure. Of note, inducing positive radial strain in diastole via modifications to the leaflet geometry – increased belly curvature and coaptation length – has proved effective in mitigating adverse tissue remodelling in TEHVs, indicating the role of tissue strain response in its remodelling behaviour^10^. Native heart valve tissue prevents the above-mentioned pathological cycle through its complex mechanical properties, evolved to minimize circumferential-to-radial strain anisotropy and maintain ECM homeostasis under hemodynamic pressure.

Native heart valve tissue is highly anisotropic with considerably lower stiffness in the radial direction^23,24^. This facilitates radial stretch in response to hemodynamic pressure and less strain anisotropy than with greater radial stiffness. Heart valve tissue stress-strain behaviour is also highly non-linear, approximately exponential^23,24^. An important property of this exponential relationship is that stress is linearly related to stiffness; therefore, cells regulate ECM stiffness by directly sensing and regulating stress present within the ECM components^21^. Despite the known importance of heart valve tissue biaxial mechanical properties, due to their complexity, until recently these properties had not been fully replicated in TEHVs^25^.

In a recent study, we reported the design, optimization, and fabrication of polymeric scaffolds composed of stacked layers of orthogonally oriented sinusoidal polycaprolactone fibres with highly tunable biaxial mechanical properties^25^. Fabricated using melt electrowriting (MEW), these scaffolds were shown to accurately and fully mimic the nonlinear, anisotropic stress-strain behaviour of native soft connective tissues with a wide range of biaxial mechanical properties. In the present study, pediatric pulmonary valves (PVs) were engineered from hybrid tissues composed of such polymeric scaffolds and cell-laden fibrin hydrogel (**Figure 1**). The architecture of the polymeric scaffolds was tuned to create four groups of pulmonary valves with native or non- native mechanical properties in the radial and circumferential direction of the valve in a combinatorial manner. This setup enabled testing the hypothesis that tissue-engineered PVs with biomimetic biaxial mechanical properties mitigate hallmarks of maladaptive tissue remodelling – including cellular differentiation to contractile phenotypes, fibrosis, and reorientation of cells toward the circumferential direction – in vitro, compared to valves with non-physiological mechanics. Testing these engineered PVs in a pulse duplicator bioreactor under physiological pulmonary pressures revealed that replicating native tissue radial mechanics significantly improved their hemodynamic performance, reduced contractile cell activation, better preserved cell orientation, and avoided tissue fibrosis. This contrasted valves that replicated native tissue mechanics only in the circumferential direction, which had poorer hemodynamic performance and evidence of fibrotic tissue remodelling. This study substantiates the role of tissue mechanical properties in directing healthy or pathological cell and ECM remodelling in TEHVs and proposes a strategy to mitigate adverse tissue remodelling by recapitulating native biaxial mechanical properties, hence creating physiological strain response under hemodynamic pressures.

**Figure 1.**
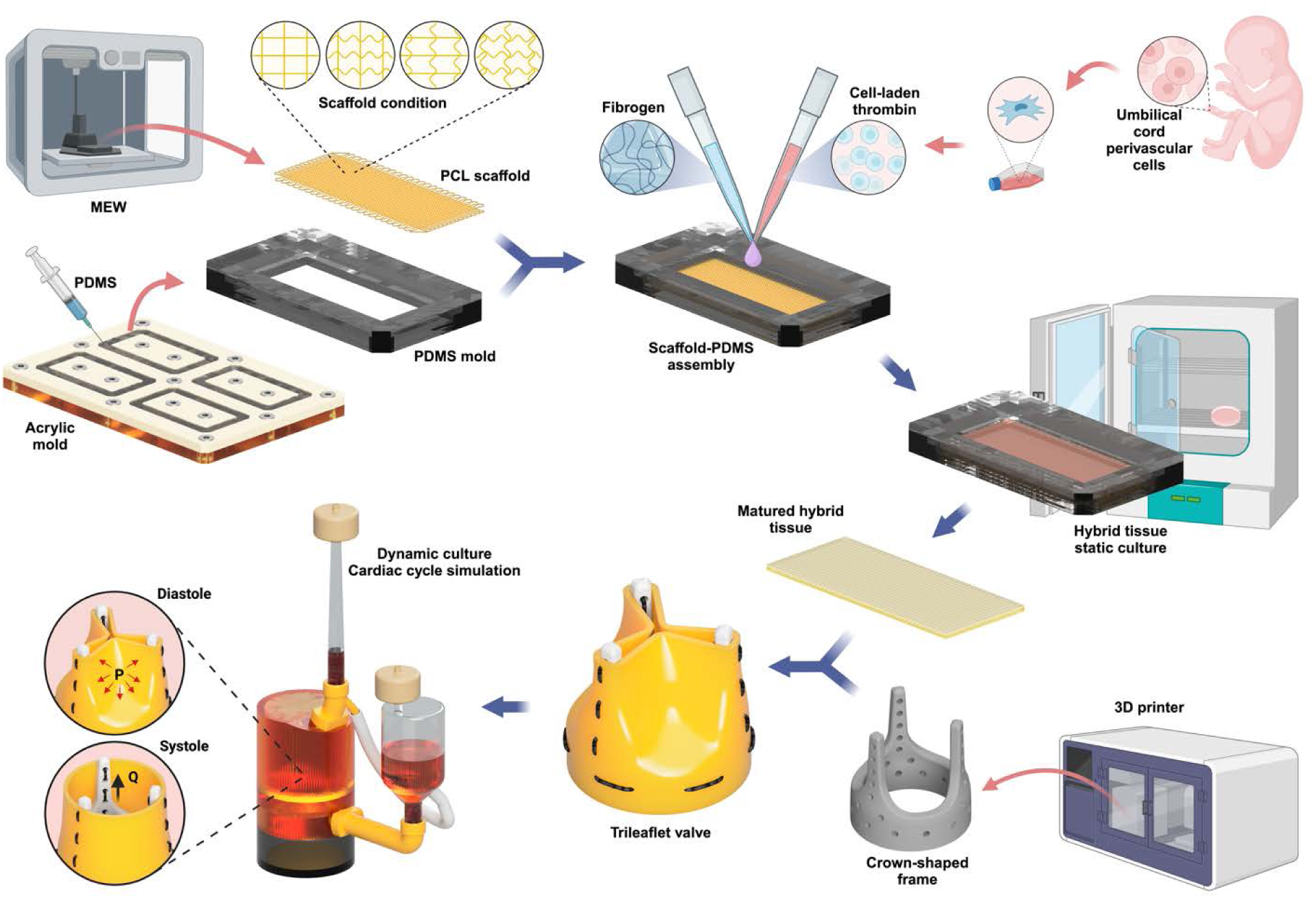
Generating tissue-engineered pulmonary valves composed of hybrid tissues with combinatorial native or non-native radial and circumferential biaxial mechanical properties and evaluating their performance and remodelling in vitro. Melt electrowriting (MEW) of polycaprolactone (PCL) was used to fabricate fibrous scaffolds with four different fibre architectures. The scaffolds were attached to polydimethylsiloxane (PDMS) molds and cellularized using fibrin hydrogel as a cell carrier (**Figure S1**). After maturation under static culture, the hybrid tissue sheets were transformed into trileaflet valves and tested under pulmonary hemodynamic conditions in a pulse duplicator bioreactor. Created with BioRender.com.

## 2- Results

### Characterization of engineered PV tissue sheets with biomimetic biaxial mechanical properties

Tissue-engineered pediatric PVs were comprised of 15 x 40 mm^2^ tissue sheets, which were statically matured from hybrid structures composed of melt electro-written polycaprolactone scaffolds and fibrin hydrogel laden with human umbilical cord perivascular cells (hUCPVCs). Using a full factorial design of experiments, the concentration/density of the constituents of the cell-laden hydrogel – including fibrinogen, thrombin, and cells – was optimized to minimize the contraction of the hybrid structure during culture (**Figure S2**) and reduce the impact of tissue culture on polymeric scaffold architecture and, therefore, hybrid tissue mechanical properties. In our preliminary studies using the optimized cell-laden hydrogel composition, hybrid tissue sheets that were matured in static culture conditions for three weeks and then transformed into trileaflet heart valves exhibited homeostatic valvular function under hemodynamic loading for five days (equivalent to 0.5 × 10^6^ cycles) in a pulse duplicator bioreactor (**Figure S3**). In contrast, tissue engineered valves constructed from sheets matured for one or two weeks failed within two days in the bioreactor.

To understand the cellular and extracellular changes that made the three-week matured hybrid tissues hemodynamically competent, cell viability and proliferation, contractile cell activation, and ECM composition throughout the static culture period were characterized. Cell viability was 82 ± 8% immediately after seeding the hybrid tissue sheets but increased significantly to 91% ± 2% by one week and to 92% ± 3% at the end of the three week culture period (p<0.03 Week 0 vs. other weeks; **Figures 2A-i** **and ii**). Cell density also increased significantly from (1.0 ± 0.2) × 10^6^ cells/ml initially to (4.9 ± 0.2) × 10^6^ cells/ml after three weeks of tissue culture (p<0.0001 Week 0 vs. other weeks; **Figure 2A-iii**). Immediately after seeding, 67% ± 5% of cells were positive for 𝛼- smooth muscle actin (SMA), consistent with the majority of cells being activated to myofibroblasts because of prior expansion of cells on a hard substrate (polystyrene flasks) substrate^26^. However, by one week of culture in soft fibrin hydrogels, the proportion of 𝛼-SMA-positive cells decreased significantly to 26% ± 7% and remained similarly low afterwards (p<0.003 Week 0 vs. other weeks; **Figure 2B**).

**Figure 2.**
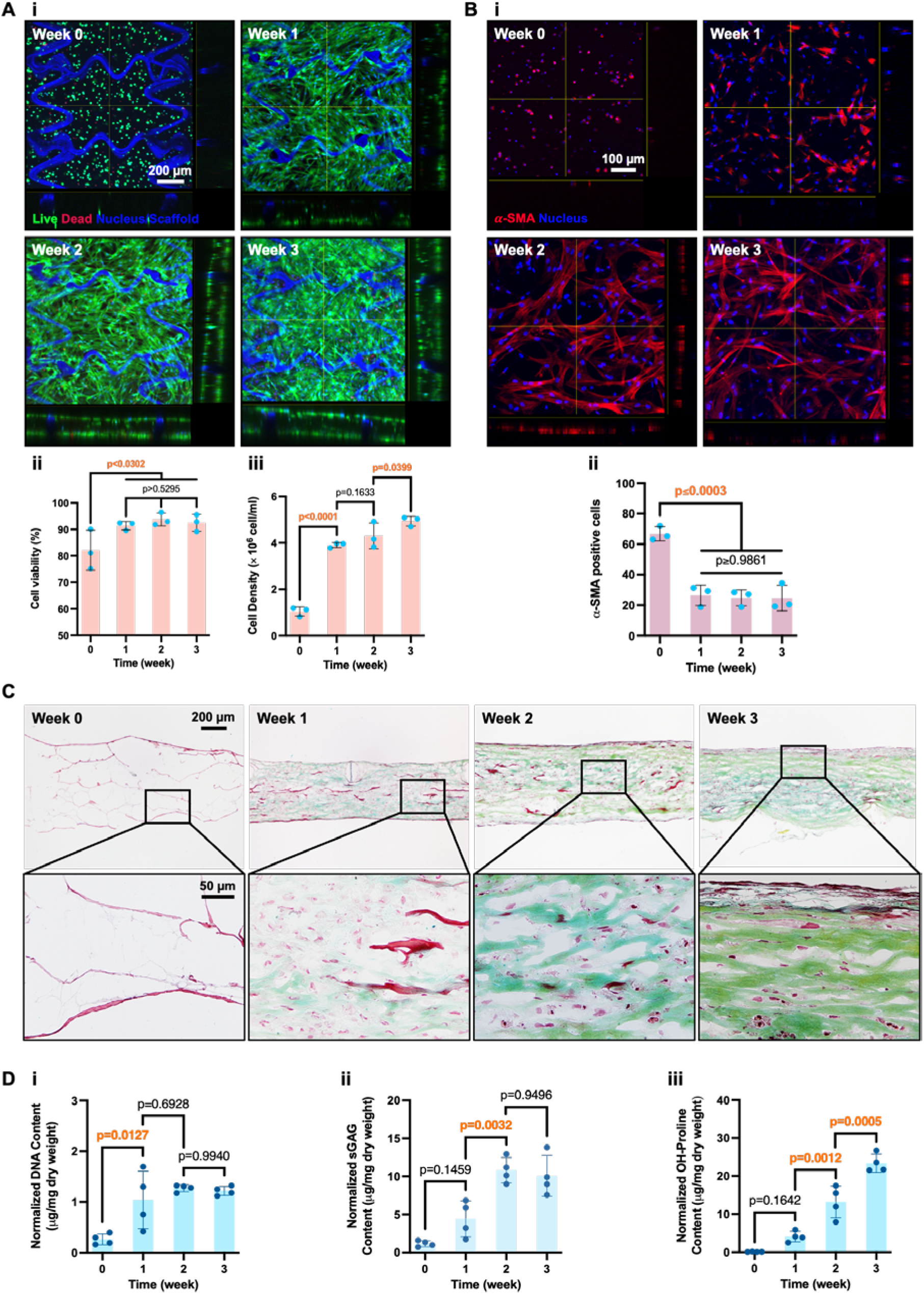
A) Laser confocal microscopic images visualizing hybrid tissues stained by Live/Dead kit (i). Cell viability (ii) and density (iii) were determined over three weeks of tissue culture. **B)** Laser confocal microscopic images of hybrid tissues stained for 𝛼-smooth muscle actin (SMA) and nuclei using Hoechst (i), and quantified for 𝛼-SMA-positive cells (ii). **C)** Microscopic images showing the cross-section of hybrid tissues stained with Movat’s pentachrome. **D)** DNA (i), sulfated glycosaminoglycan (sGAG) (ii), and collagen (iii) contents of hybrid tissues were quantified using biochemical analysis and normalized to dry tissue weight. Data are presented as mean ± SD with N=3 (A, B) and N=4 (D).

Histological analysis using Movat’s pentachrome staining showed gradual degradation of fibrin hydrogel (bright red), deposition of glycosaminoglycans (blue) mainly between week one and two, and deposition of collagen (yellow) predominantly after week two (**Figure 2C**). Biochemical analysis showed an increase in DNA content, especially during the first week of tissue maturation (**Figure 2D-i**), in agreement with the Live/Dead assays (**Figure 2A-iii**). Biochemical assays also showed an increase in sulfated glycosaminoglycan content up to week two of culture (**Figure 2D- ii**) and an increase in collagen content throughout the course of tissue maturation (**Figure 2D-iii**), both consistent with histological staining (**Figure 2C**).

The accumulation and maturation of ECM over three weeks’ culture putatively contributed to the mechanical integrity of the hybrid tissues, enabling them to withstand hemodynamic loading (**Figure S3**). We hypothesized that the ECM synthesized over three weeks would also affect the hybrid tissue’s biaxial mechanical properties, causing them to deviate from those of the native pediatric PV tissue that the polymeric scaffold was designed to emulate. Indeed, whereas the first version of the polymeric scaffold (PV_V1_), developed in our previous study^25^, (**Figure 3Ai**) matched the biaxial mechanical properties of native PV within 5 ± 2% (**Figure 3Aiii**), three weeks of tissue formation (**Figure 3Aii**) caused the hybrid tissue sheet to become less extensible and stiffen in both the circumferential and radial directions (**Figure 3Aiii**), deviating by up to 38% from native mechanics (**Figure 3Bi**). From the difference between the tension-strain behaviours of the bare polymer scaffold and three-week matured hybrid construct, the contribution of the cell-laden hydrogel was calculated and used to derive, via linear regression (**Figure 3B**), scaffold architectural parameters that would yield biomimetic biaxial tension-strain behaviour in the resultant three-week matured hybrid tissues. Polymeric scaffolds with the optimized architecture (PV_V2_) were melt-electrowritten, cultured for three weeks as before, and tested mechanically, at which point their tension-strain behaviour was similar to that of native tissue, with just 9.4% ± 4.5% and 9.6% ± 4.7% deviation in the radial and circumferential directions, respectively (**Figure 3C**).

**Figure 3.**
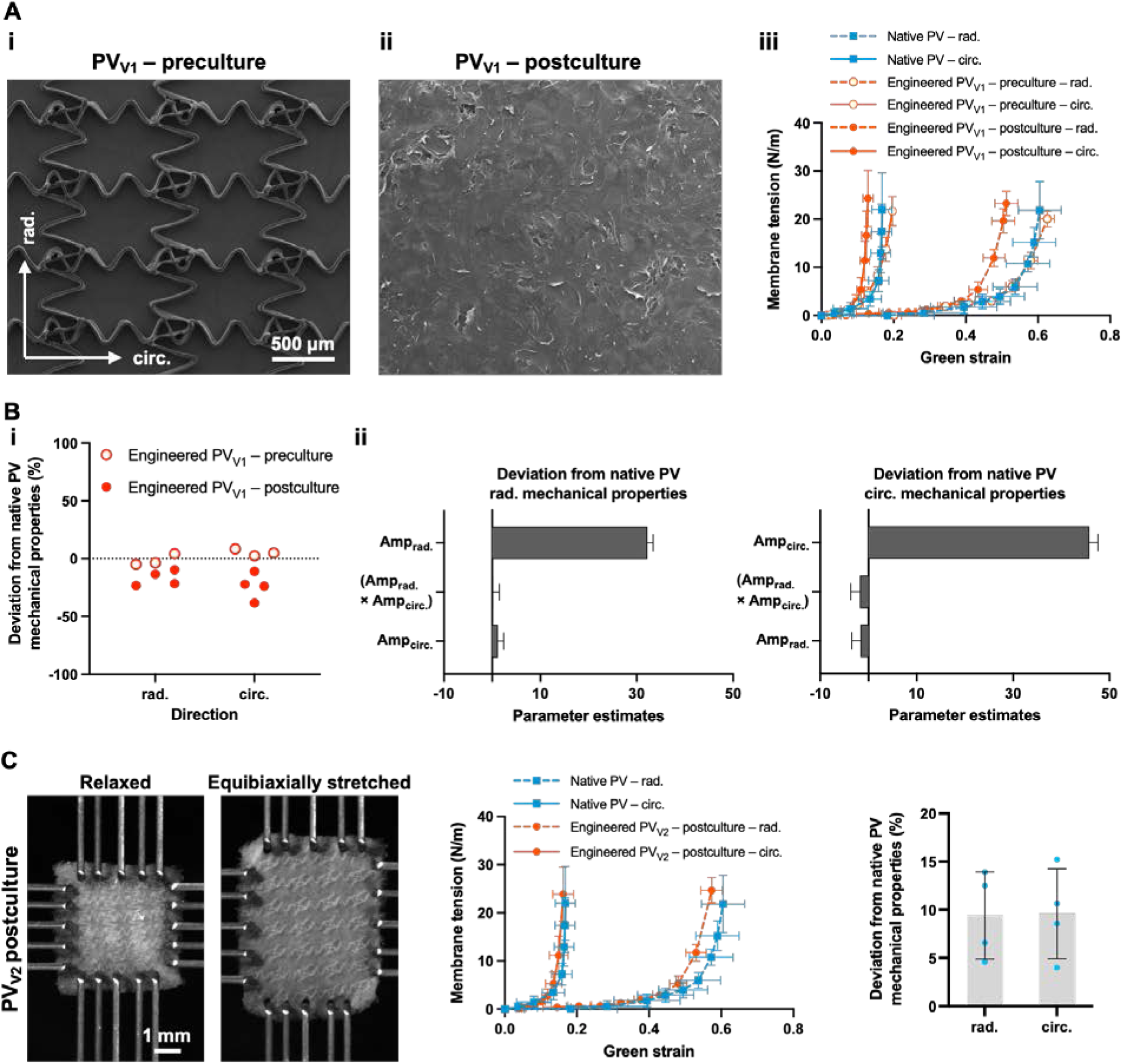
A) Polymeric scaffold previously developed using melt electrowriting that was shown to recapitulate native pulmonary valve (PV) tissue biaxial mechanical properties (i). A hybrid structure composed of the polymeric scaffold and cell-laden fibrin hydrogel, cultured for three weeks (ii). The biaxial mechanical properties of the polymeric scaffold, hybrid tissue, and native PV tissue (iii). **B)** The deviation of polymeric scaffold and hybrid tissue mechanical properties from those of native tissue in the radial and circumferential direction (i). The relationship between the polymeric scaffold fibre wave amplitude (Amp) and the deviation of corresponding scaffolds’ mechanics from that of native tissue was determined using a regression model (ii). **C)** The regression model was used to optimize the architectural parameters of the polymeric scaffold to yield native-like mechanical properties. The hybrid tissue composed of the optimized polymeric scaffold underwent biaxial mechanical testing and showed tension-strain behaviour similar to that of native tissue. Data are presented as mean ± SD with N=4.

### Development of engineered PVs with combinatorial biaxial mechanical properties

Four types of engineered PVs were defined to determine the impact of replicating native biaxial mechanical properties in TEHVs on their hemodynamic performance, cellular behaviour, and ECM remodelling. These engineered PVs possessed combinatorial native or non-native mechanical properties in radial and circumferential directions (**Figure 4A**). We used a two-digit identifier for the four conditions, where the first number represents the radial direction and the second number represents the circumferential direction, and with values of “0” representing non-native mechanics and “1” representing native mechanics. Following this nomenclature, the four conditions tested were: PV00 with non-native mechanics in both radial and circumferential directions; PV01 with non-native mechanics in radial but native mechanics in the circumferential direction; PV10 with native mechanics in radial but non-native mechanics in circumferential direction; and PV11 with native mechanical properties in both radial and circumferential directions. In these engineered PVs, native mechanics was provided by optimized sinusoidal fibre architecture and non-native mechanics was achieved by straight fibres **(Figure 4B)**, anticipating straight fibres would generate tissue stiffness higher than native levels. The non-native conditions mimic the majority of current tissue-engineered heart valves tested in vivo, which show stress-strain behaviour with stiffness higher than that of native tissue^5,6,27,28^. In particular, PV01 with similar tension-strain behaviour in the radial and circumferential direction mechanically emulated TEHVs with isotropic mechanics, such as those developed from nonwoven polymeric sheets with random fibre orientation and tested in vivo^10^.

**Figure 4.**
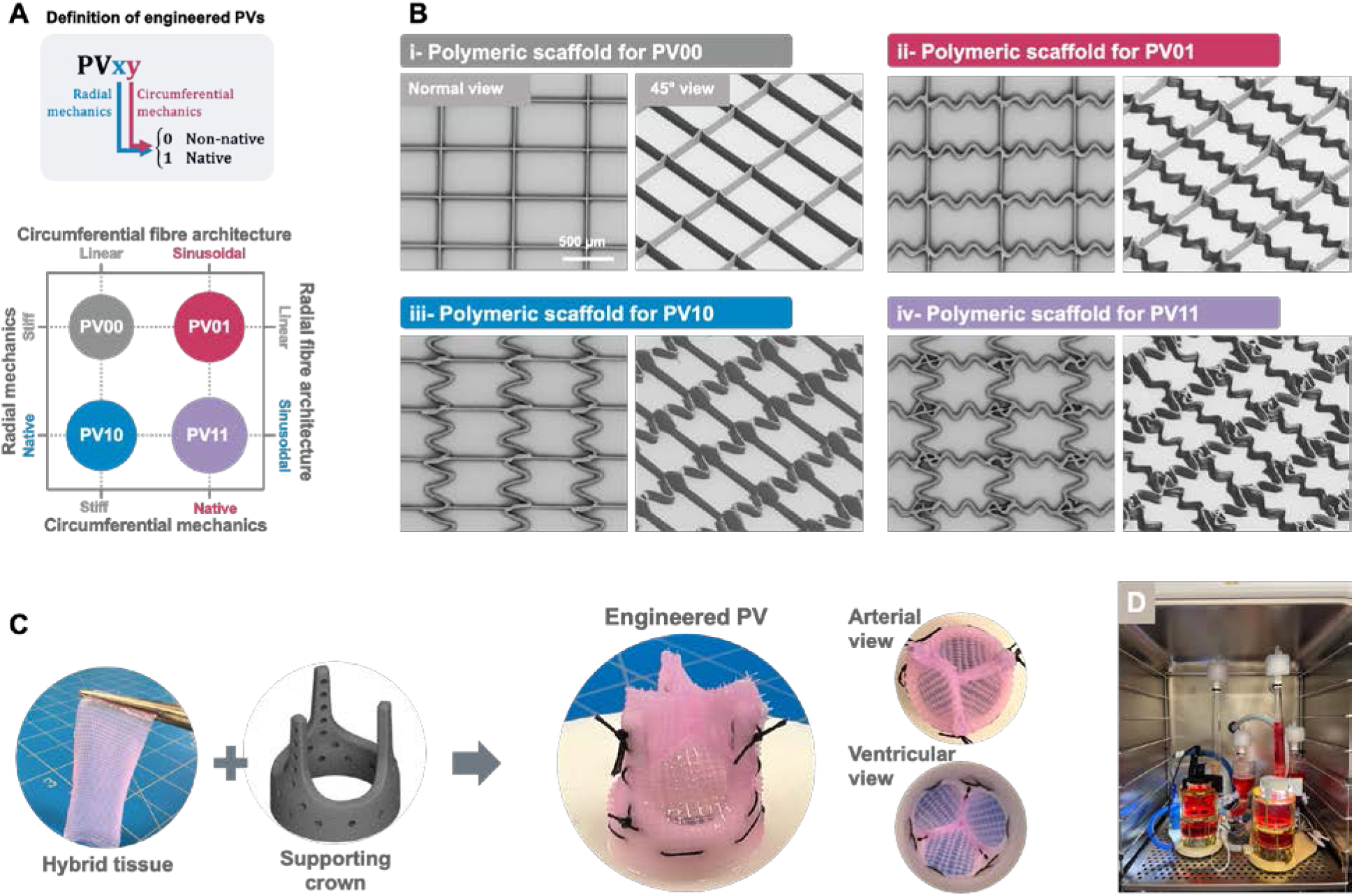
A) Definition of four types of engineered pulmonary valves (PVs) with combinatorial native or non-native mechanical properties in radial and circumferential directions: PV00 with non-native mechanics in both radial and circumferential directions; PV01 with non-native mechanics in radial but native mechanics in circumferential direction; PV10 with native mechanics in radial but non-native mechanics in circumferential direction; and PV11 with native mechanical properties in both radial and circumferential directions. **B)** Scanning electron micrographs of four types of polymeric scaffolds used to engineer the four types of PVs described above. **C)** Fabrication process of trileaflet PVs using hybrid tissue sheets, composed of polymeric scaffolds and cell-laden fibrin hydrogel, and crown-shaped supporting frames.

### Cell viability and proliferation of engineered PVs

The four types of hybrid tissue sheets – composed of polymeric scaffolds (**Figure 4B**) and cell-laden hydrogel, matured for three weeks – were transformed into trileaflet engineered PVs using crown-shaped supporting frames (**Figure 4C**) and cultured dynamically under pulmonary conditions in a pulse duplicator bioreactor (**Figure 4D**). Construction of trileaflet valves required aseptic handling of the hybrid tissue under room atmospheric conditions with non-ideal temperature, humidity, and CO_2_ concentration to maintain cell viability. With an initial design of the crown-shaped frame, the valve assembly process took ∼50 min and led to significant cell death, reducing the cell viability from 88% ± 7% to 22% ± 7% (p<0.0001) (**Figure S4**). A modification on the crown design reduced the valve assembly time to ∼25 min and increased the cell viability to 45% ± 5% (p=0.0090) (**Figures S4, 5A** and **B-i**).

Despite this cell loss during the valve assembly process, cell viability increased significantly during the 5-day dynamic culture period to 69% ± 6% (p=0.0005) (**Figures 5B-i** and **C**), reaching live cell densities comparable to those in the original tissue sheets before the trileaflet valve was constructed (p=0.065) (**Figure 5B-ii**). The recovery of cell viability and density was attributable to proliferation rates during bioreactor culture ((0.29 ± 0.06) × 10^6^ cells/ml/day) that were comparable to those in static culture (**Figure 5D**). No significant difference was observed between the four types of engineered PVs in cell viability (p>0.87) and cell density (p>0.12) (**Figure 5E**). Of note, the dead cell nuclei after dynamic culture were smaller than those observed before and after the trileaflet valve assembly process (**Figure S5**). These likely represented remnants of cells that died during valve assembly, and since a portion were included in the post-bioreactor viability measurement, the calculated viability of 69% ± 6% underestimated the actual viability. The significant increase in cell viability and density during hemodynamic function revealed the capability of all four types of engineered PVs to recover from the detrimental effect of the valve assembly process, while maintaining valvular function.

**Figure 5.**
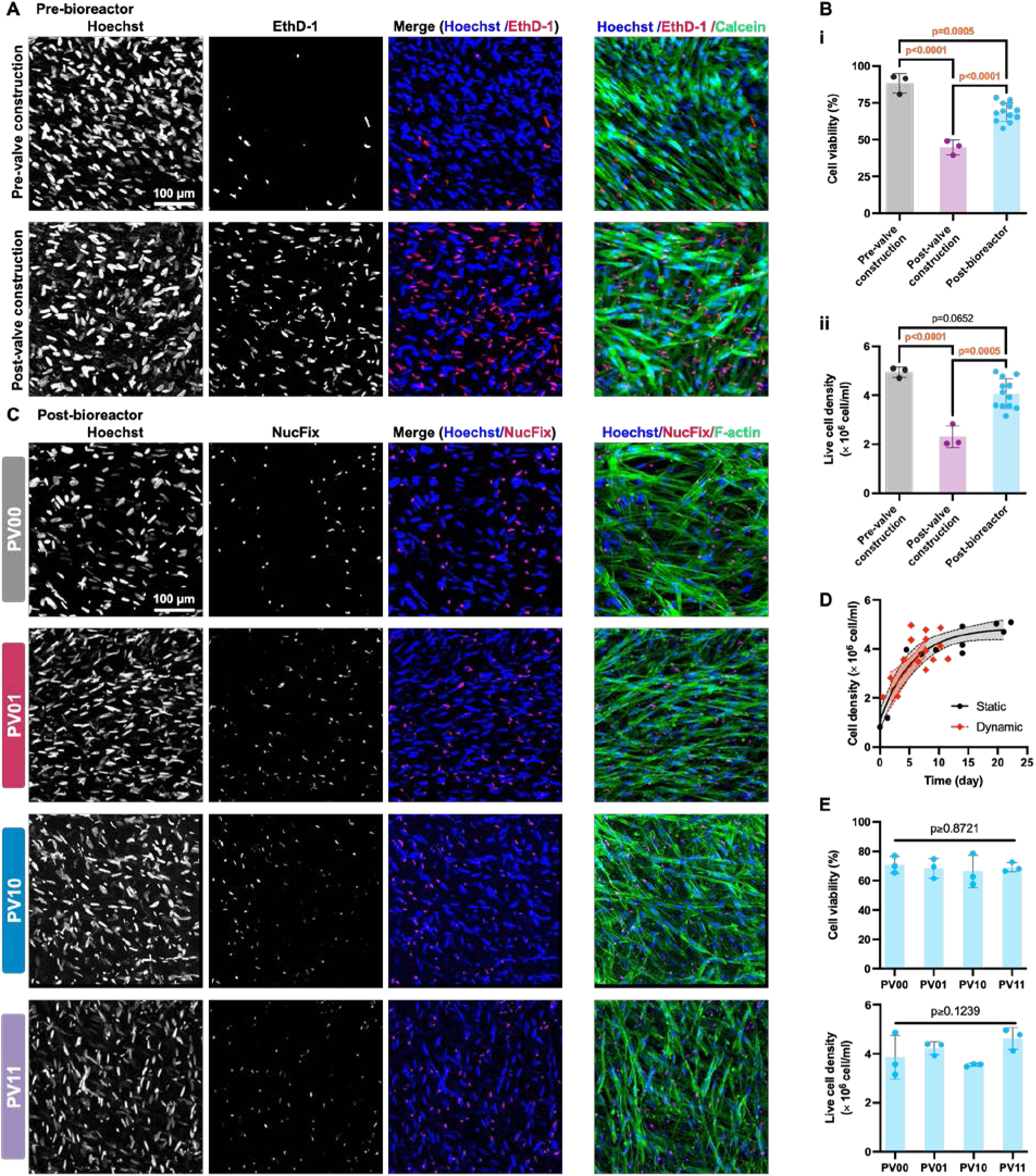
A) Laser confocal microscopic images of hybrid tissues stained with Hoechst (all cell nuclei), ethidium homodimer-1 (EthD-1) (dead cell nuclei), and Calcein (live cell cytoplasm), showing the cell viability before and after transformation of hybrid tissues into trileaflet pulmonary valves (PVs). **B)** Cell viability (i) and density (ii) in engineered PV tissues before and after sheet-to-trileaflet transformation and after hemodynamic function for 5 days (0.5 × 10^6^ cycles). **C)** Laser confocal microscopic images of engineered PV leaflets, stained with Hoechst, NucFix (dead cell nuclei), and phalloidin (F-actin filaments), after hemodynamic function for 5 days. PV00 with non-native mechanics in both radial and circumferential directions; PV01 with non-native mechanics in radial but native mechanics in circumferential direction; PV10 with native mechanics in radial but non-native mechanics in circumferential direction; and PV11 with native mechanical properties in both radial and circumferential directions. **D)** Cell density increased similarly during static and dynamic culture. **E)** Cell viability and density in the four types of engineered PVs after the hemodynamic function. Data represent mean ± SD with N=3 for each group of hybrid tissue and engineered PVs.

### Biaxial mechanical properties of engineered PVs

The tension-strain behaviour of hybrid tissues after three weeks of static culture, before transformation into trileaflet valves, was determined via biaxial tensile testing. As anticipated, straight fibres generated stiffness higher than native levels, and fibres with optimized sinusoidal morphology generated tension-strain behaviours that matched those of native PV tissue (**Figure 6**). As a result, the four types of hybrid tissues used for the fabrication of engineered PVs showed combinatorial native or stiff (i.e., stiffer than native levels) radial and circumferential mechanical properties: PV00 with straight fibres in radial and circumferential directions had stiff tension-strain behaviour in both directions (**Figure 6A**); PV01 with straight and sinusoidal fibres in the radial and circumferential direction had stiff and native mechanics in respective directions (**Figure 6B)**; PV10 with sinusoidal and straight fibres in the radial and circumferential direction, had native and stiff mechanics in respective directions (**Figure 6C)**; and PV11 with sinusoidal fibres in both directions had full native biaxial tension-strain behaviour (**Figure 6D**). The four types of PVs were loaded in the bioreactor and underwent hemodynamic loading for 5 days under pulmonary conditions, after which their leaflet tissues were retrieved and tested for their biaxial mechanical properties. Among the four types of valves, PV00 and PV01, both with non-native radial mechanics, showed changes in mechanical properties after hemodynamic loading. The extensibility of PV00 increased in both directions (**Figure 6A**), and the extensibility of PV01 increased only in the radial direction, the direction in which the valve did not emulate native mechanical properties (**Figure 6B**). This was likely due to the breakage of the polymer fibres, putatively due to the rigidity of the scaffold in the radial (in PV00 and PV01) and circumferential (in PV00) direction (**Figures S6A** and **B**). Additionally, differential scanning calorimetry showed an increase in the crystallinity of polycaprolactone from 45% ± 2% to 51% ±45 (p=0.0025) during valvular function in PV00 (**Figure S7**), which is indicative of strain-induced crystallinity, making the scaffold susceptible to damage. In contrast, there was no evidence of damage to the polymeric scaffold fibres in PV10 and PV11, the scaffolds with native radial mechanics (**Figures S6C** and **D**). Consistent with this, the mechanical properties of PV10 and PV11 also did not change after 5 days of hemodynamic loading (**Figures 6C** and **D**).

**Figure 6.**
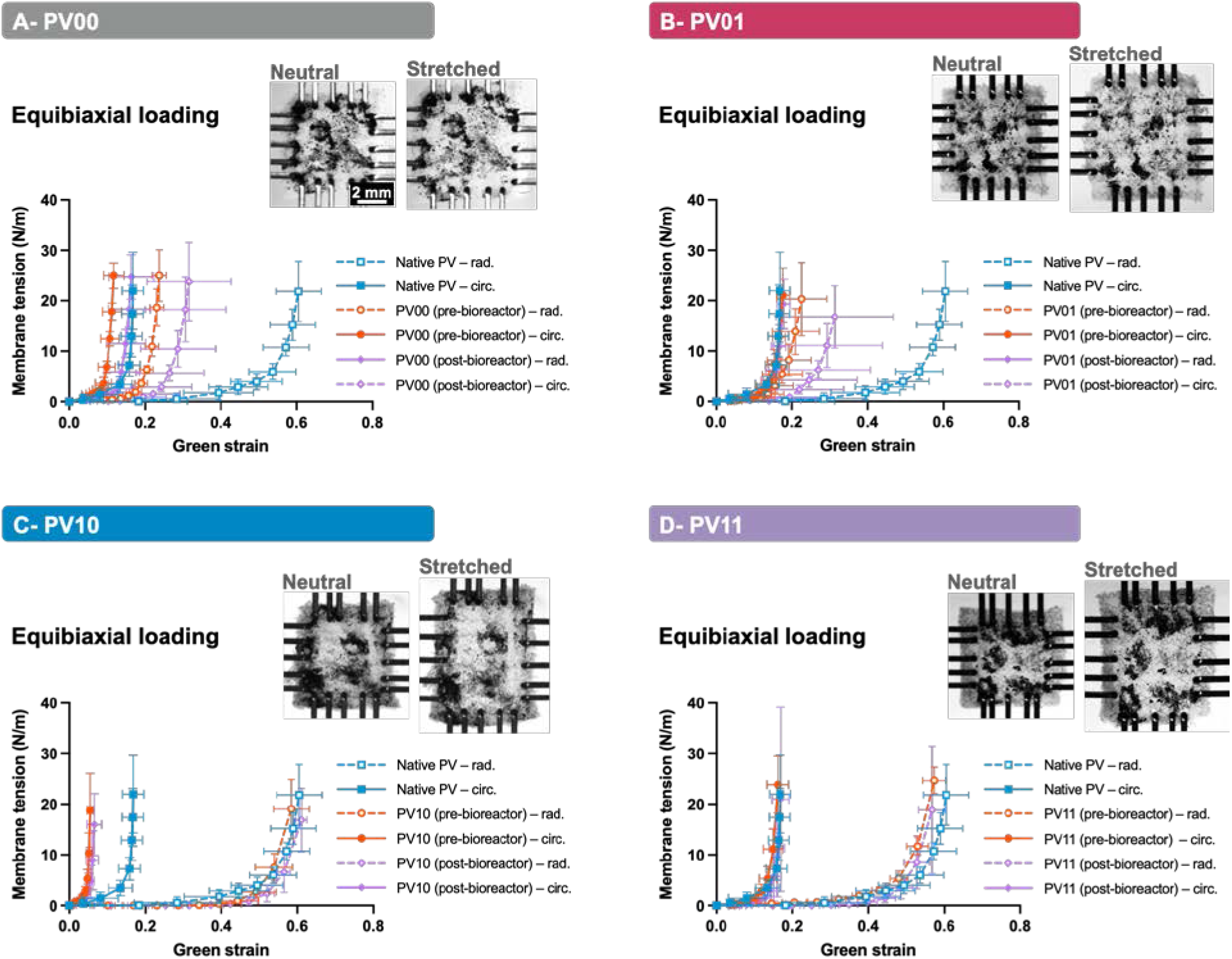
Biaxial membrane tension-strain relationship of four types of pulmonary valves (PVs) – PV00 (**A**), PV01 (**B**), PV10 (**C**), and PV11 (**D**) – before and after hemodynamic loading and valvular function for 5 days (pre- and post- bioreactor) in comparison with biaxial mechanical properties of native pediatric PV tissue^24^. Data are presented as mean ± SD with N=4.

### Hemodynamic performance of engineered PVs

Under the pulmonary pressure and flow rate in a pulse duplicator bioreactor, engineered PVs with non-native radial mechanics, PV00 and PV01, showed incomplete diastolic closure (**Figures 7A- I and ii**); in contrast, valves with native radial mechanics, PV10 and PV11, closed completely in diastole (**Figures 7A-iii and iv**). This was evidenced in both high frame-rate imaging during the valvular function and the pressure profiles, in which PVs with native radial mechanics exhibited more stable diastolic pressure profiles (**Figure 7A**). Determined by image analysis and post-processing of pressure profiles, PV10 and PV11 had smaller diastolic orifice areas (p≤0.0027) (**Figure 7B-i**) and greater diastolic transvalvular pressures (p≤0.0343) (**Figures 7B-ii and iii**) than PV00 and PV01, the valves with non-native radial mechanics, throughout the 5-day hemodynamic function period. Among the four types of PVs, only PV00 showed variable diastolic transvalvular pressure (**Figure 7B-ii**), presumably owing to the damage to its polymeric scaffold that allowed better closure over time. Nonetheless, diastolic transvalvular pressure in PV00 remained significantly lower than PV10 and PV11 and marginally lower than PV01 throughout the dynamic culture period. In systole, the valves with native radial mechanics (PV10 and PV11) consistently had lower transvalvular pressures than PV00 and PV01 (p≤0.019) (**Figures 7C-i**, and **ii**), indicating lower resistance to flow. This can be explained by the lower radial bending resistance observed in hybrid tissues with native radial mechanics compared to their stiff counterparts (**Figure S8**). Of note, no significant differences in hemodynamic function were found between the two PVs with native radial mechanics (PV10 and PV11) and between the two with non-native radial mechanics (PV00 and PV01).

**Figure 7.**
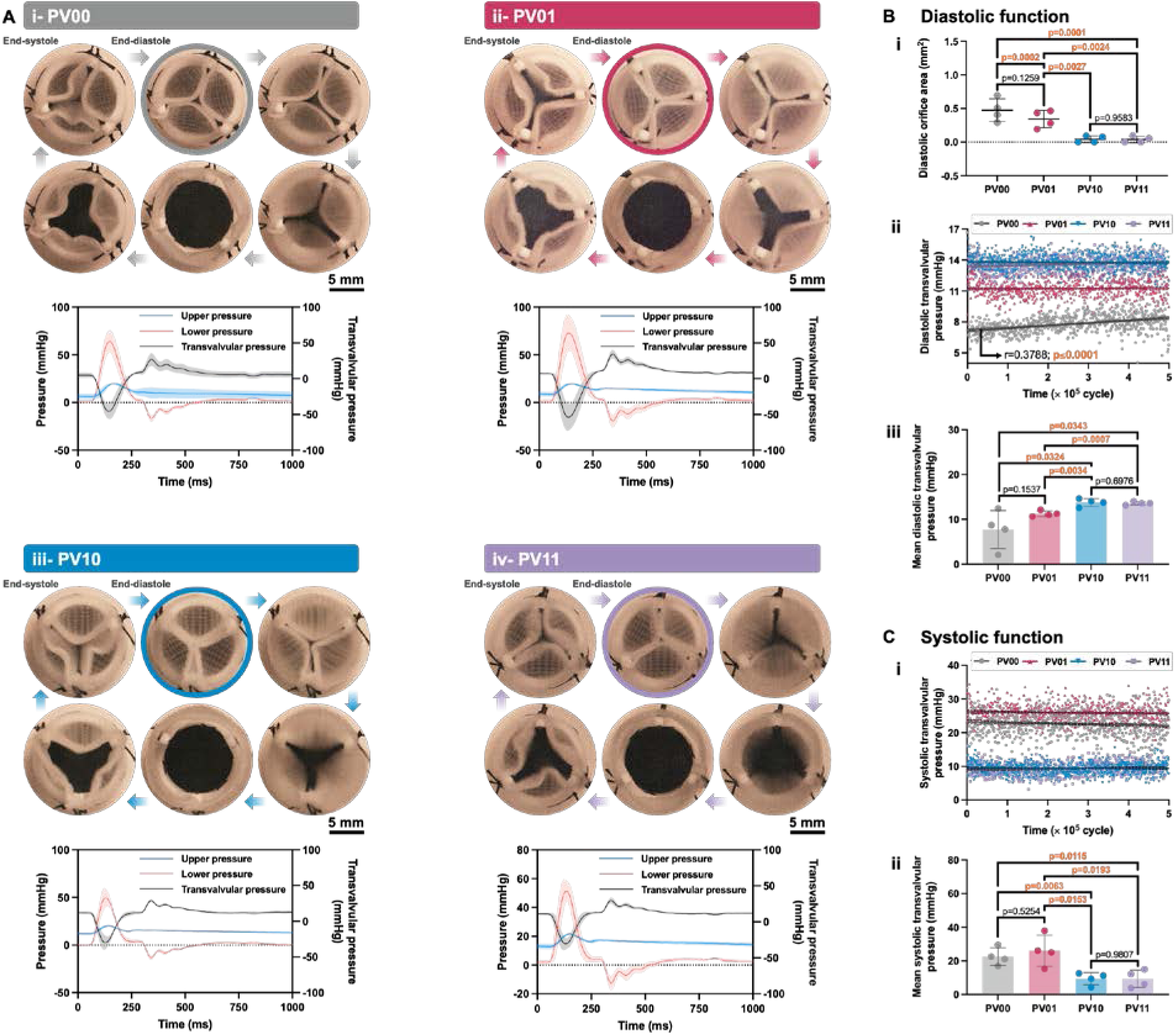
A) Engineered pulmonary valves (PVs) in one cardiac cycle: PV00 with non-native mechanics in both radial and circumferential directions (i); PV01 with native mechanics in circumferential but non-native mechanics in radial direction (ii); PV10 with native mechanics in radial but non-native mechanics in circumferential direction (iii); and PV11 with native mechanical properties in both radial and circumferential directions (iv). In the graphs, solid lines and shaded areas show mean ± SD. **B)** Diastolic function of the four types of engineered PVs including diastolic orifice area (i), diastolic transvalvular pressure over 0.5 × 10^6^ cycles (ii), and mean diastolic transvalvular pressure (iii). **C)** Systolic function of the four types of engineered PVs including systolic transvalvular pressure over 0.5 × 10^6^ cycles (i) and mean systolic transvalvular pressure (ii). Data are presented as mean ± SD with N=4 for each PV type.

Considering native PV hemodynamic characteristics^29,30^, engineered PVs with native radial mechanics showed normal physiological diastolic and systolic performance well far from diseased conditions (regurgitation and stenosis); however, PV00 showed marginally healthy diastolic and systolic function and PV01 showed marginally healthy systolic function (**Figure S9**).

### Contractile cell activation in engineered PVs

Leaflet retraction and fibrotic remodelling in TEHVs *in vivo* have been shown to associate with activation of contractile cells, primarily shown by their expression of 𝛼-SMA^2,5,10^. This has been shown in TEHVs containing multiple cell types, such as dermal fibroblasts^5,31^, bone marrow- derived stem cells^32,33^, and peripheral vein-derived fibroblasts^34^, as well as in decellularized^5,6,10^ or cell-free^12,27^ TEHVs that were cellularized endogenously *in vivo*. As a hallmark of tissue fibrotic remodelling, 𝛼-SMA expression by hUCPVCs in the engineered PVs after dynamic culture was investigated. To achieve a complete map of 𝛼-SMA expression, one leaflet of each engineered PV underwent immunostaining and a complete laser confocal microscopic scan (**Figure 8A**). The results showed that recapitulating native radial mechanics in engineered PVs (PV10 and PV11) significantly reduced contractile cell activation under hemodynamic loading compared to PVs with non-native radial mechanics (**Figure 8B**). While PV00 and PV01 showed no significant difference in expressing 𝛼-SMA (p=0.8672), their 𝛼-SMA expression was 4.4-fold higher than engineered PVs with native radial mechanics, PV10 and PV11 (p<0.0073) (**Figure 8C**). Comparing 𝛼-SMA expression at the end of the static to the dynamic culture, engineered PVs with native radial mechanics maintained the cell phenotype during the dynamic culture (p≥0.7168). In contrast, PVs with stiff radial mechanics had increased differentiation to the contractile phenotype (p≤0.0001) (**Figure 8C**).

**Figure 8.**
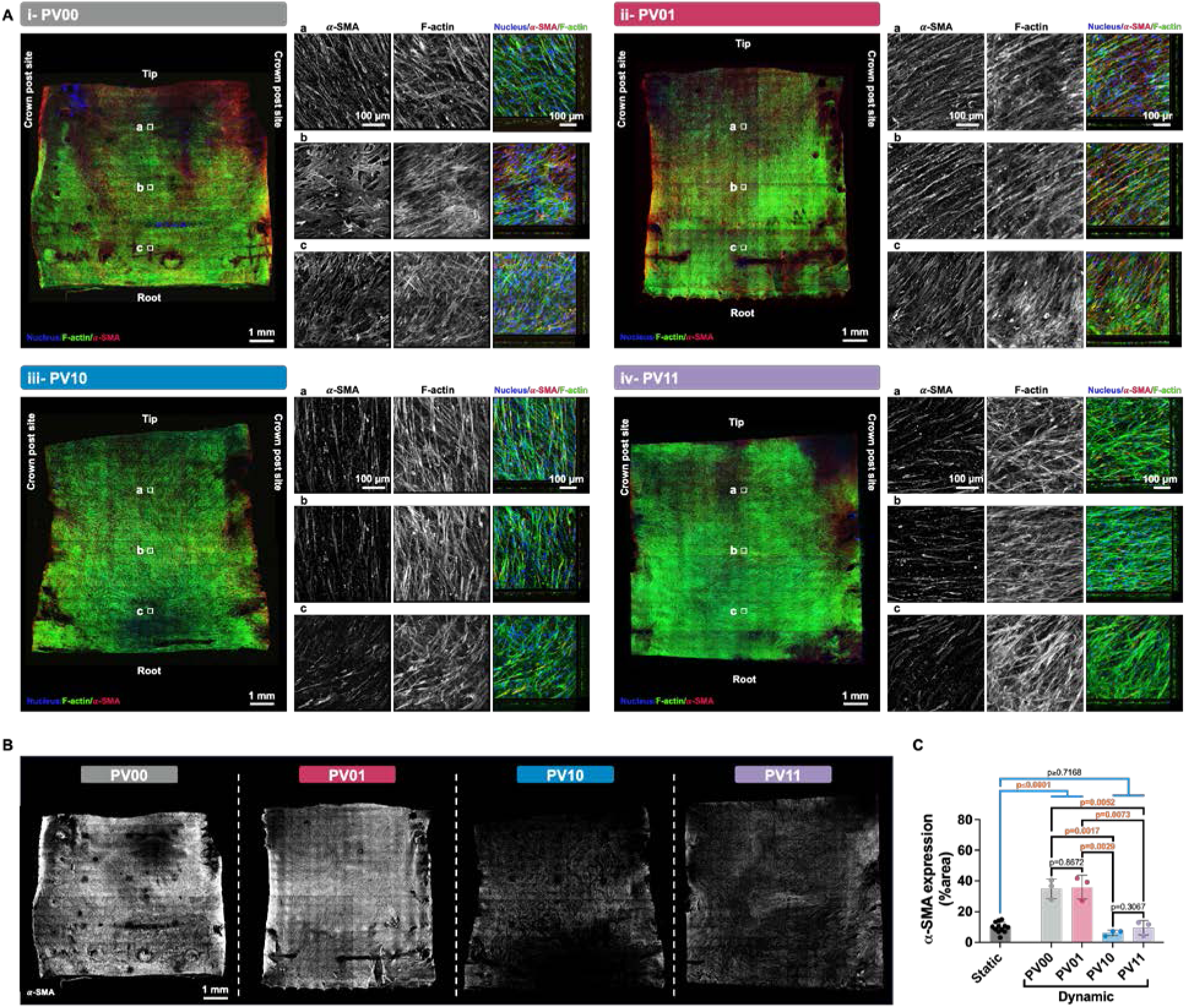
A) Laser confocal microscopic images developed by scanning whole leaflets of four types of engineered pulmonary valves (PVs) stained with Hoechst for nuclei (blue), phalloidin for F-actin (green), and anti 𝛼-smooth muscle actin (SMA) to identify myofibroblasts (red). PV00: non-native mechanics in both radial and circumferential directions; PV01: non-native mechanics in radial but native mechanics in circumferential direction; PV10: native mechanics in radial but non-native mechanics in circumferential direction; PV11: native mechanical properties in both radial and circumferential directions. **B)** The expression of 𝛼-SMA in the four types of engineered PVs. **C)** Proportion of the leaflet area with positive 𝛼-SMA expression for each of the four types of engineered PVs. Data are presented as mean ± SD with N=3 for each PV type.

### Cell orientation in engineered PVs

Reorientation of cells toward the circumferential direction has been shown as another hallmark of tissue retraction and fibrotic remodelling in TEHVs^18^. Derived from computational modelling, this phenomenon has been attributed to non-physiological strain anisotropy caused by negligible or negative strain in the radial direction of the valve leaflet tissue^18,20^. Therefore, to assess the potential of fibrotic tissue remodelling in the four types of engineered PVs, the orientation of cells in the entirety of one leaflet retrieved from each valve was determined by the quantification of F- actin filament orientation and compared to the orientation of cells in hybrid tissues prior to hemodynamic loading. Prior to valvular function, cells were randomly oriented in all four types of hybrid tissues (**Figure 9A-i**), with no preferential orientation (**Figure 9A-ii**). After 5 days of hemodynamic loading, cells throughout much of the PV leaflets with non-native radial mechanics oriented more circumferentially, with 33% ± 5% and 35% ± 7% of cells circumferentially oriented in PV00 and PV01, respectively (**Figures 9B** and **C**). PV10 leaflets showed a reduced trend in cell reorientation compared to PV00 and PV01, with 28% ± 4% of cells oriented toward the circumferential direction. In contrast, significantly fewer cells were circumferentially oriented in PV11 leaflets with native radial and circumferential mechanics (22% ± 2%; p=0.024 vs. PV00, p=0.035 vs. PV01). Importantly, unlike all other PV conditions, PV11 with full native mechanics had no significant change in circumferential orientation of cells after 5 days of hemodynamic loading compared to hybrid tissues at the end of static tissue maturation (19% ± 3%; p=0.18).

**Figure 9.**
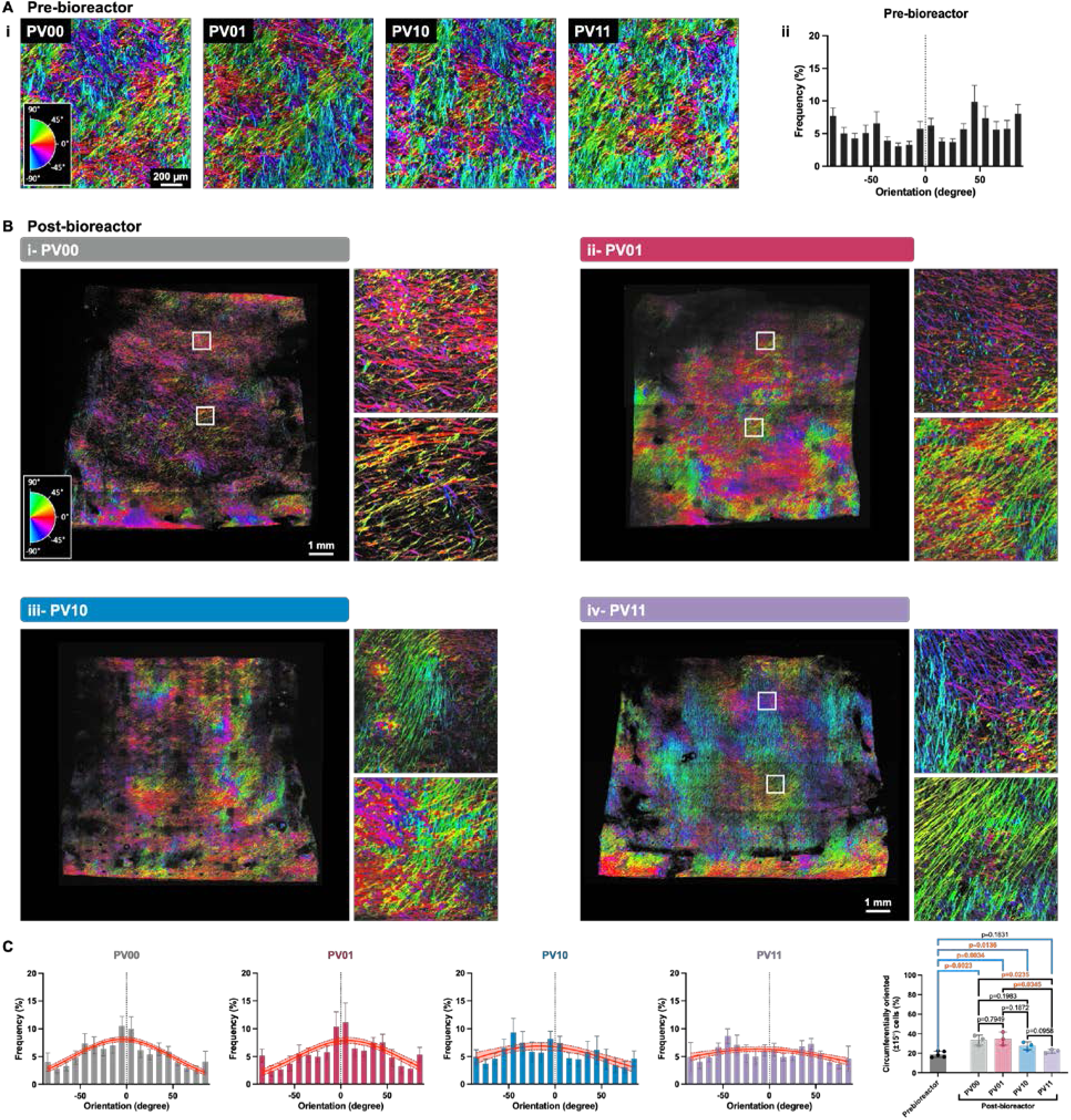
A) Pseudo-coloured laser confocal microscopic images (i) and distribution of cell orientation (ii) in four types of hybrid tissues after static tissue maturation and before the construction of pulmonary valves (PVs). PV00: non-native mechanics in both radial and circumferential directions; PV01: native mechanics in circumferential but non-native mechanics in radial direction; PV10: native mechanics in radial but non-native mechanics in circumferential direction; PV11: native mechanical properties in both radial and circumferential directions. **B)** Pseudo-coloured laser confocal microscopic images of engineered PVs, with combinatorial radial and circumferential mechanical properties, after valvular function. **C)** Distribution of cell orientation in the four types of engineered PVs after valvular function. Data are presented as mean ± SD with N=3 for each PV type.

### Collagen expression in engineered PVs

Leaflet retraction in TEHVs associates with excessive deposition of collagen^2^. Therefore, as a hallmark of tissue fibrotic response, the expression of collagen in the four types of engineered PVs was determined after 5 days of hemodynamic loading using multiphoton microscopy and compared to that of hybrid tissues at the end of static culture (**Figures 10A** and **B**). Collagen levels were comparable between the four PV types prior to dynamic culture, with 25% ± 5% of the leaflet areas positive for collagen fibres (**Figures 10A** and **C**). After dynamic culture, there was a significant increase in collagen in PVs with non-native radial mechanics (41% ± 2% and 47% ± 7% for PV00 and PV01, respectively) (**Figures 10B-i, B-ii, and C**; p≤0.0002 vs. pre- bioreactor). In contrast, after dynamic loading, the amounts of collagen in PV10 (30% ± 4%) and PV11 (29 ± 4%) with native radial mechanics were not different from the tissues prior to dynamic loading (p=0.19) and were significantly lower than in PV00 and PV01 (p≤0.019) (**Figures 10B- iii, B-iv, and C**). These results indicate the potential for fibrosis in PVs with non-native, stiff radial mechanical behaviour, and the ability of PVs with highly extensible, native radial tension-strain behaviour to promote ECM homeostasis. In contrast, replicating native circumferential mechanical properties had no effect on avoiding tissue fibrosis and maintaining tissue collagen content.

**Figure 10.**
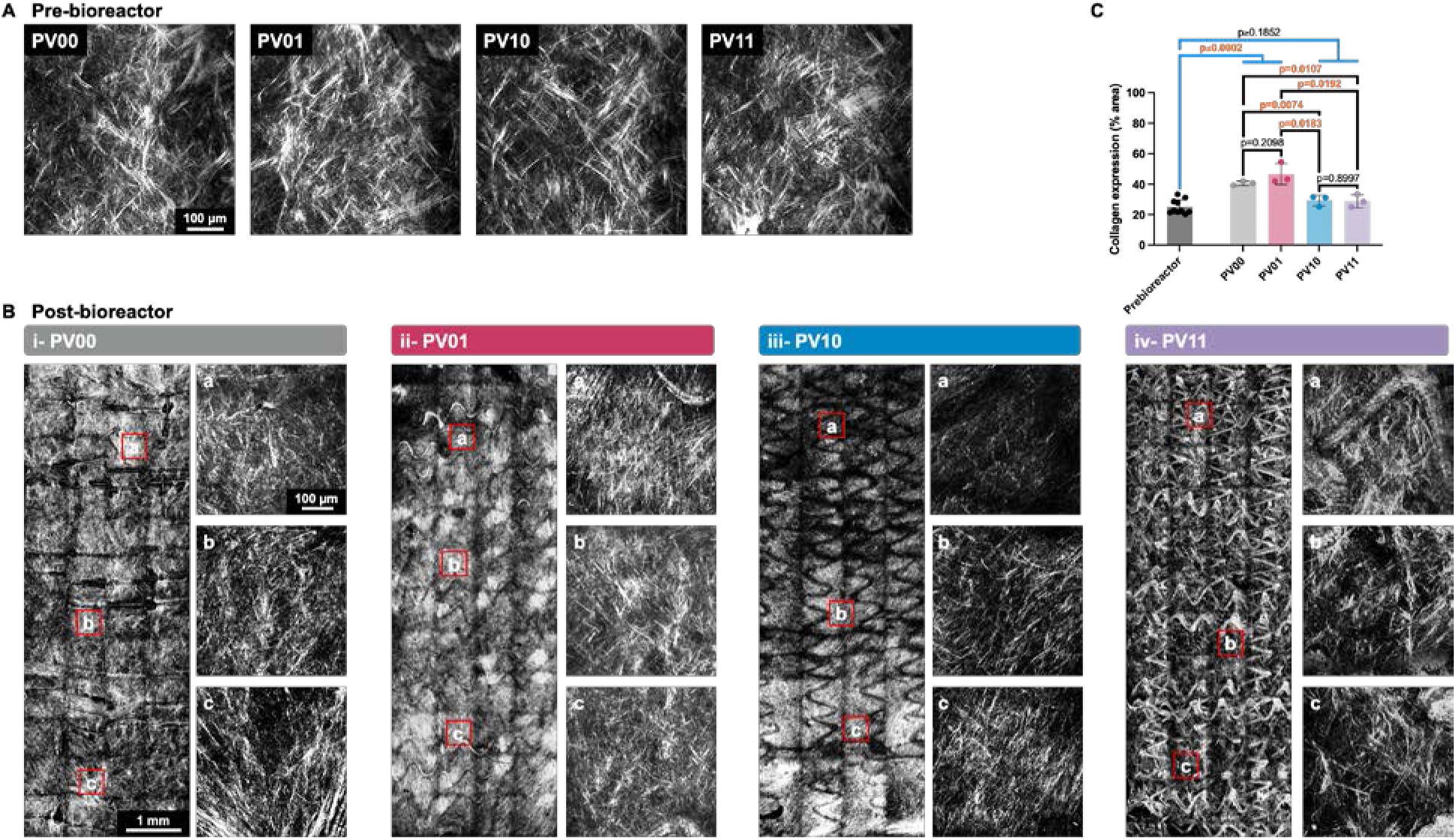
A) Multiphoton microscopic images showing collagen fibres in four types of hybrid tissues after static tissue maturation and before the construction of pulmonary valves (PVs). PV00: non-native mechanics in both radial and circumferential directions; PV01: non-native mechanics in radial but native mechanics in circumferential direction; PV10: native mechanics in radial but non-native mechanics in circumferential direction; PV11: native mechanical properties in both radial and circumferential directions. **B)** Multiphoton microscopic images showing collagen fibres in engineered PVs, with combinatorial radial and circumferential mechanical properties, after valvular function. **C)** Collagen expression in the four types of engineered PV tissues before and after valvular function. Data are presented as mean ± SD with N=3 for each PV type.

### Histological analysis

Movat’s pentachrome staining of hybrid tissues after three weeks of static culture (**Figure 11A**) and engineered PV leaflets’ radial sections (**Figure 11B**) after 5 days of valvular function confirmed engineered PVs with non-native, stiff radial mechanics remodel fibrotically under dynamic loading, whereas PVs with highly extensible, native radial tension-strain behaviour promote ECM homeostasis. This was evident by more smooth muscle cells (dark red in **Figures 11B-i** and **ii**) and more collagen (yellow in **Figures 11B-i** and **ii**) in PV00 and PV01 than in PV10 and PV11 (**Figures 11B-iii** and **iv**), consistent with the 𝛼-SMA immunostaining (**Figure 8**) and collagen imaging (**Figure 10**) data.

**Figure 11.**
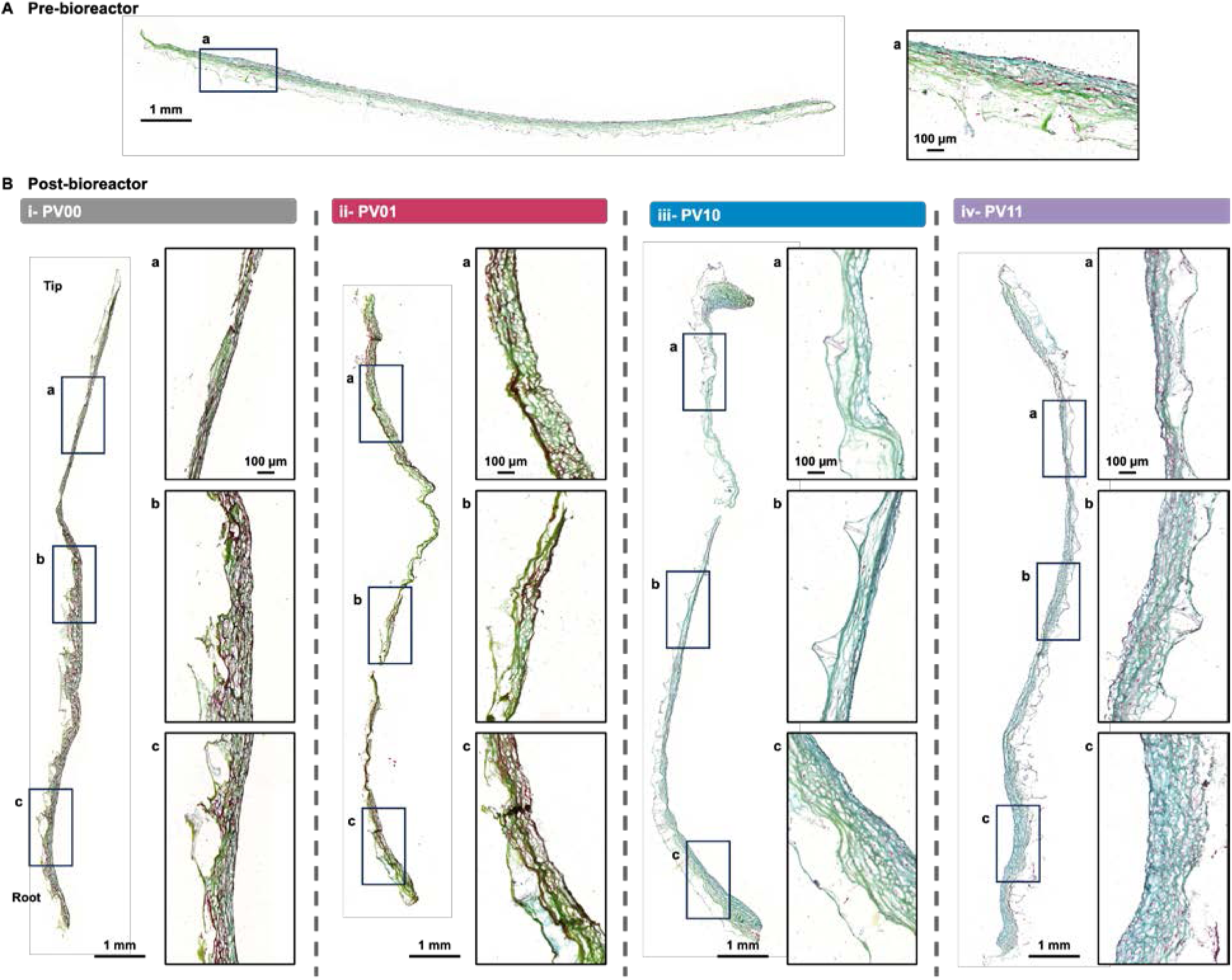
A) Microscopic image of a hybrid tissue matured for three weeks stained with Movat’s pentachrome. **B)** Microscopic images of engineered pulmonary valve (PV) leaflets’ radial section stained with Movat’s pentachrome. PV00 (i) with non-native mechanics in both radial and circumferential directions; PV01 (ii) with non-native mechanics in radial but native mechanics in circumferential direction; PV10 (iii) with native mechanics in radial but non-native mechanics in circumferential direction; and PV11 (iv) with native mechanical properties in both radial and circumferential directions.

### Engineered PVs strain response to diastolic pressure

Computational^18^ and experimental^22^ studies suggest a critical role for strain response on heart valve tissue remodelling behaviour – cell contractility and reorientation and fibrosis. Therefore, a testing device was developed to simulate the diastolic phase of the cardiac cycle and expose engineered PVs to physiological diastolic transvalvular pressure under micro-computed tomography (𝜇CT) imaging (**Figure S10**). 𝜇CT scan enabled the reconstruction of the engineered PV geometry in neutral and pressurized conditions (**Figure 12A**), as well as visualizing radial and axial sections of the valve leaflets (**Figure 12B**) to calculate leaflet belly curvature and strain in the radial and circumferential directions. As anticipated, valves with native radial mechanics had larger belly curvatures (**Figure 12** **C**) and radial strain (13% ± 3% and 12% ± 3% in PV10 and PV11, respectively) (**Figure 12D-i**) compared to those with stiff radial mechanics (2% ± 2% and - 1% ± 1% radial strain in PV00 and PV01, respectively). In contrast, valves with stiff radial mechanics experienced trivial or negative radial strains, indicating reduced translation of hemodynamic pressure to radial strain. As a result, strain anisotropy was significantly higher (p≤0.0211) in engineered PVs with stiff radial mechanics (9.9 ± 3.7 and 10.9 ± 2.7 in PV00 and PV01, respectively) compared to their counterparts with native radial mechanics (2.6 ± 0.8 and 1.4 ± 0.4 in PV10 and PV11, respectively).

**Figure 12.**
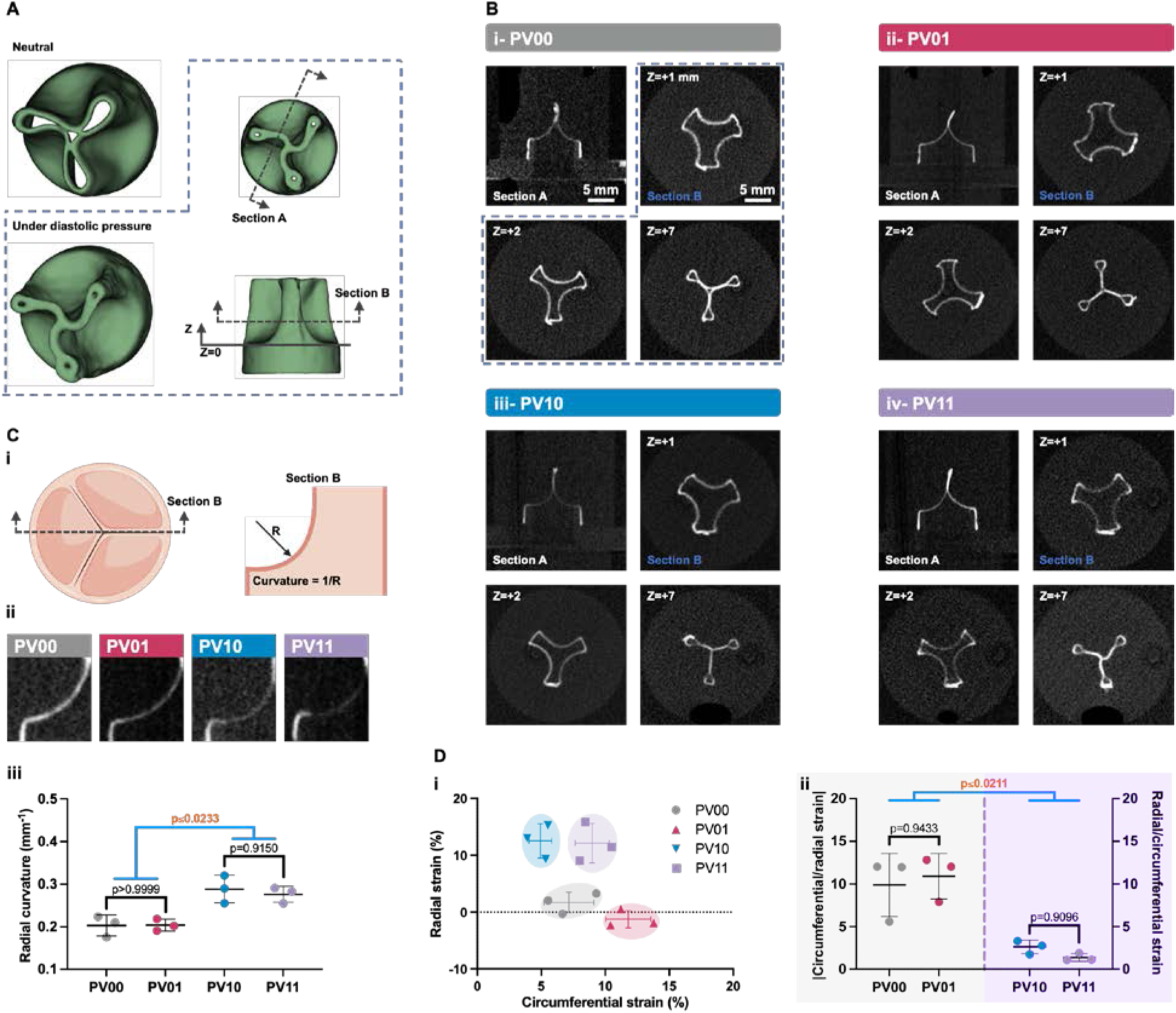
A) Constructed 3D model derived from the micro-computed tomography (𝜇CT) scans of an engineered pulmonary valve (PV) inside the diastolic test device in neutral and pressurized conditions. ***B)*** Radial and axial sections of the four types of engineered PVs under the pulmonary diastolic transvalvular pressure (10 mmHg). PV00 (i) with non-native mechanics in both radial and circumferential directions; PV01 (ii) with non-native mechanics in radial but native mechanics in circumferential direction; PV10 (iii) with native mechanics in radial but non-native mechanics in circumferential direction; and PV11 (iv) with native mechanical properties in both radial and circumferential directions. ***C)*** Schematic demonstration of leaflet belly radial curvature in engineered PVs (i). Created with BioRender.com. Axial section of 𝜇CT scans, showing the belly radial curvature in the four types of engineered PVs under the pulmonary diastolic transvalvular pressure (ii), and associated quantification (iii). ***D)*** Radial and circumferential strain (i) and strain anisotropy (ii) experienced by the four types of engineered PVs under the pulmonary diastolic transvalvular pressure at their belly region. In (ii), for each valve type, the larger strain ratio between circumferential/radial strain and radial/circumferential strain is reported. Data are presented as mean ± SD with N=3 for each PV type.s

## 3- Discussion

Causing leaflet retraction and valve regurgitation, adverse tissue remodelling in TEHVs has been characterized with the occurrence of myofibroblast activation, circumferential cell reorientation, and excessive deposition of collagen^1,6,10^. While computational modelling has been utilized to simulate trileaflet valves’ geometry, boundary conditions and hemodynamic loading in order to reveal the underlying mechanobiological causes of these processes^18^, experimental implementation of such studies with physiologically relevant conditions has been a challenge. Based on previous studies suggesting that abnormally high strain anisotropy (circumferential/radial strain) accelerates adverse tissue remodelling in TEHVs^18,20,35^, here we hypothesized that this pathological tissue remodelling can be mitigated by recapitulating native valve tissue biaxial mechanical properties. This hypothesis was tested by fabricating various types of TEHVs with native and non-native tension-strain behaviour and monitoring the hallmarks of pathological tissue remodelling under physiological hemodynamic loading.

The MEW method, by providing superior control over fibre architecture, allowed for accurate and precise prescription of stress-strain behaviour in the circumferential and radial direction of engineered heart valves. Subsequently, engineering heart valves with combinatorial biaxial mechanical properties enabled investigating the effect of TEHVs’ circumferential and radial mechanical properties, independently, on their valvular performance and tissue remodelling. The four types of PVs engineered in this study, generated a wide range of mechanical properties from native PV mechanics to non-native mechanical properties, simulating those of TEHVs previously tested *in vivo*^5,6,27,28^.

As they are under lower hemodynamic pressure during valvular function, engineered PVs are more susceptible to tissue contraction compared to engineered aortic valves (AVs)^18^. It has been shown that lower pressures present in the PV environment allow the valve tissue to experience low or negative radial stretch, which accelerates adverse tissue remodelling if its resident cells possess the potential to exhibit contractility^11,18^. Therefore, engineered PVs are a suitable platform to study the effect of their mechanical properties on their cell and ECM homeostasis.

Among the four types of engineered PVs, those with native radial mechanics (PV10 and PV11) showed no damage to the polymeric scaffold during valvular function and outperformed those with non-native radial mechanics (PV00 and PV01) in diastolic and systolic performance. PV10 and PV11 maintained higher diastolic pressure, indicative of lower backflow, and created lower resistance to flow in systole. Similar valvular performance of PV00 and PV01 (both with non- native mechanics), as well as PV10 and PV11 (both with native radial mechanics), showed that recapitulating circumferential mechanics (i.e., in PV01) is of less significance compared to the radial mechanics in enhancing the valves’ hemodynamic function.

Hybrid structures, in which the pores of the polymeric scaffold were filled with cell-laden fibrin hydrogel, suppressed contractile cell activation during static tissue maturation. The number of 𝛼- SMA^+^ cells decreased by 2.6-fold (**Figure 1B-ii**) from the beginning of tissue culture to week 1 and remained constantly low for up to three weeks. This may be attributed to the downregulation of myosin-dependent contractility that has been shown to happen in soft matrices^21,36^. While the relatively stiff polymeric scaffold governed, in large part, the mechanical properties of the hybrid tissue, the hydrogel component, and later the secreted natural ECM, shielded the cells from contractile activation, which has been shown to occur on stiff substrates in a series of events including the development of large focal adhesions, high traction force, and eventually actomyosin-mediated contractility^37^. Among four types of engineered PVs, those with native radial mechanics, PV10 and PV11, maintained cell quiescence with reduced myofibroblast activation and collagen synthesis under dynamic loading during valvular function. This was associated with generating a radial strain response, larger than the other two conditions, to the hemodynamic pressure in diastole by mimicking native tissue tension-strain behaviour in the radial direction. In the valves with non-native radial mechanics, PV00 and PV01, in which radial tissue stiffness was considerably larger than physiological levels, leaflet tissue did not experience stretch in the radial direction under pulmonary hemodynamic conditions, particularly because of the unconstrained free-edge of the valve leaflet. In contrast, hemodynamic pressure was translated to circumferential stretch given the anchored boundary conditions at the commissure sites in this direction. Together, this led the valve tissues with stiff radial mechanics to experience higher strain anisotropy in diastole than those with native radial mechanics. In a previous study, the Butcher group reported that the expression of ACTA2 mRNA, which encodes 𝛼-SMA in valvular interstitial cells, increased with strain anisotropy^22^. This is consistent with our finding, in which the high expression of 𝛼-SMA in PV00 and PV01 was associated with the larger strain anisotropy in these valves under hemodynamic pressure.

The large radial stiffness in engineered PVs with non-native radial mechanics caused the leaflet tissue in PV00 and PV01 to experience trivial and negative strain in the radial direction, respectively, which impacted cell reorientation^18^. It has been shown that under biaxial strain, cells avoid orientation toward the direction of compaction and orient toward the direction in which they experience positive strain^20^. This is because actin filaments are more stabilized in the direction of tension, as tension reduces actin depolymerization rate^38^ and its severing by cofilin^39^. This phenomenon can explain the larger degree of circumferential reorientation in PV00 and PV01, as these valves had the highest radial stiffness and hence the largest potential to experience negative radial strain. in contrast, PV11 with native mechanical properties and large extensibility in the radial direction preserved cell orientation at the highest degree among the four types of PVs.

Overall, our data show that recapitulating native tissue mechanics in TEHVs, particularly in the radial direction, supports valvular performance and tissue homeostasis. TEHVs with non-native, stiff radial mechanics showed hallmarks of adverse fibrotic tissue remodelling, including contractile cell activation, increased collagen synthesis, and reorientation of cells toward the circumferential direction. In contrast, engineered PVs with native non-linear, anisotropic mechanical properties maintained cell quiescence, preserved cell orientation, and suppressed collagen synthesis, all indicative of tissue homeostasis. This work, by providing strong evidence on the superior performance of TEHVs with native nonlinear, anisotropic mechanical properties, motivates future investigation to assess whether such TEHVs mitigate adverse tissue remodelling, seen in current TEHVs, in the long term *in vivo*.

## 4- Conclusion

The impact of radial and circumferential mechanical properties in TEHVs on their valvular performance and tissue remodelling was studied via engineering four types of PVs with combinatorial native or non-native radial and circumferential mechanical properties. Engineered PVs with native radial mechanics, with or without native circumferential mechanics, showed superior valvular performance in diastole and systole. Such valves also showed minimal contractile cell activation and fibrosis. In contrast, replicating only native circumferential mechanical properties in engineered PVs did not improve their valvular function. Such valves showed signs of adverse tissue remodelling, including contractile cell activation, circumferential cell reorientation, and increased collagen synthesis. The results of this study showed that recapitulating native heart valve tissue mechanical properties, particularly in the radial direction, in TEHVs is essential to promote their cell and ECM homeostasis. Knowing the issue of *in vivo* tissue retraction in current TEHVs has remained a challenge, this study suggests a practical approach to mitigate this adverse tissue remodelling and pave the way toward clinical translation of TEHVs.

## 5- Methods

### Fabrication of scaffolds

A glass syringe (Cadence Science, Italy, cat# G5117, 5 mL) was filled with 1 g of polycaprolactone pellets (Sigma-Aldrich, UK, cat# 704105; Mn = 42 kDa, Tg = −63 °C, and Tm = 55.7 °C, as characterized previously^25^), kept in an oven at 90 °C for 8 h, and stored in a dark, dry chamber at room temperature for future use. A stainless steel, 32-gauge needle (Hamilton, USA, cat# 7747-02 10 mm PT3) was attached to the syringe, and the assembly was inserted into a custom-made MEW machine with the syringe and needle heated to 90°C and 95°C, respectively. TCL code was used to control the movement of the MEW machine in X-Y-Z directions. Scaffolds composed of 10 layers of fibres in each orthogonal direction were deposited on glass microscope slides (VWR, Germany, cat# 16004-368) while voltage, print-head pressure, and working distance between the needle and the print substrate were kept at 1650 V, 225 kPa, and 0.8 mm, respectively. MEW was performed above the critical translation speed at 2 mm/s.

#### SEM

MEW scaffolds were sputter-coated (Polaron Range SC7640, Quorum Technologies, UK) with 5 nm of gold and imaged using a Prisma E SEM (ThermoFisher Scientific, USA) with a 10- kV electron beam and a secondary electron detector.

### Tissue culture

MEW scaffolds were attached to polydimethylsiloxane (PDMS) molds (as described previously^25^), sterilized using 70% ethanol, and plasma treated for two min under ambient conditions. Human umbilical cord perivascular cells (provided in-kind by Tissue Regeneration Therapeutics) at passage three were thawed and suspended at 2×10^6^ cell/ml into a complete xeno-free StemMacs culture medium (Miltenyi Biotec, USA, cat# 130-104-182) containing 1 U/ml of human thrombin (Sigma-Aldrich, USA, cat# T6884). Equal volumes of the cell suspension and a human fibrinogen (Sigma-Aldrich, USA - cat# F3879) solution (10 mg/ml) in Dulbecco’s phosphate-buffered saline (DPBS)−/− (Gibco, USA, cat# 14190-144) were cast in the PDMS mold cavity within the polymeric scaffold. Following incubation at 37 °C, 100% humidity, and 5% CO_2_ for 90 min for complete cross-linking, cell culture medium supplemented with 1% antibiotics solution (a mixture of 10 mg/ml penicillin (BioShop, Canada, cat#: PEN333.25), 10% gentamicin (Sigma-Aldrich, USA, cat#: G1397), and 30 𝜇g/ml amphotericin B (Sigma-Aldrich, USA, cat#: A9528)) was added to the resultant hybrid structure. The hybrid structure was cultured for up to three weeks, and culture media was refreshed every three days.

### Tissue staining and laser confocal microscopy

At different time points, tissues were stained by Hoechst 33342 (ThemoFisher Scientific, USA, cat # 62249) and a Live/Dead kit (Invitrogen, USA, cat# L3224), as previously described^25^. A separate set of samples was stained to visualize dead nuclei, actin filament, 𝛼-SMA, and total nuclei. In brief, tissues were stained with 1X NucFix^TM^ Red dead cell kit (Biotium, USA, cat# 32010) for 30 min, fixed with 4% paraformaldehyde in DPBS for 15 minutes at room temperature, permeabilized using 0.2% Triton X-100 (Sigma-Aldrich, USA, cat# T8532) for 10 min at room temperature, and blocked with 1% bovine serum albumin (Sigma-Aldrich, USA, cat# A9418) and 22.52 mg/mL glycine (Sigma-Aldrich, USA, cat# G8898) in PBST (DPBS + 0.1% Tween 20 (Sigma-Aldrich, USA, cat# P1379)) for 1h at room temperature. Samples were incubated in a solution containing 1:1000 anti-𝛼-SMA antibody (Abcam, UK, cat# ab202296), 1:20 phalloidin (Sigma-Aldrich, USA, cat# P5282), and 1% BSA in PBST overnight at 4°C. Samples were stained with Hoechst 33342 solution for 5 min at room temperature and kept in Aqua-Mount® mounting medium (ThermoFisher Scientific, USA, cat# CA41799-008). All stained tissues were imaged using a laser confocal microscope (Olympus FV3000, Japan). Microscopic images were analyzed using the 3D Object Counter module of Fiji software to determine cell viability using the equation below.

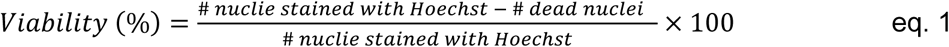

Full scans were performed on one leaflet of each engineered PV, stained with NucFix^TM^ Red, anti-𝛼-SMA, phalloidin, and Hoechst 33342, to quantify 𝛼-SMA expression and cell orientation. Each leaflet was placed in a well of a 6-well glass bottom plate (Cellvis, USA, cat# P06-1.5H-N) and sandwiched using a coverslip (Chemglass Life Sciences, USA, cat# CLS-1760-025) while suspended in Aqua-Mount® mounting medium. Leaflets were imaged with four laser channels at peak excitation of 405 nm, 490 nm, 520 nm, and 650 nm. Samples were tile-scanned using a 20X objective lens with a 3-𝜇m z-step size to cover their full thickness and area (12×12 mm^2^). The expression of 𝛼-SMA was quantified by measuring the fraction of 𝛼-SMA^+^ area of the leaflet after thresholding the images using the Otsu algorithm in Fiji. The actin filament orientation was determined using the OrientationJ plugin of Fiji.

### Multiphoton microscopy

Engineered tissue leaflets were fixed with 4% paraformaldehyde in DPBS for 15 minutes at room temperature and stored in DPBS+/+ (Corning, USA, cat# 21-030- CV) at 4°C before imaging. Samples were sandwiched between two glass coverslips, placed on the microscope (Nikon A1R MP+ with 25X Apo LWD Water Dipping Objective, Japan) stage and imaged using 930 nm two-photon exposure. Samples were tile-scanned with a 7-𝜇m z-step size to cover their full thickness and an area of 3×12 mm^2^, a radial strip at the centre of each leaflet.

### Biochemical analysis

Quantitative biochemical assays were performed to measure DNA, OH-proline, and sGAG in hybrid tissues. Post culture, tissues were washed with DPBS+/+, placed in pre-weighed centrifuge vials, flash-frozen in liquid nitrogen, and freeze-dried. Each sample was digested in 250 µL of papain digestion buffer containing 35 mM ammonium acetate (Sigma-Aldrich, USA, cat# A7330), 1 mM ethylenediaminetetraacetic acid (EDTA) (BioShop, Canada, cat# EDT111), 2 mM DL-dithiothreitol (Sigma-Aldrich, USA, cat# D9779), and 80 μg/mL papain (Sigma-Aldrich, USA, cat# P3125) at pH 6.2 on a heat block at 65°C for 72 hours. For the OH-proline assay, acid hydrolysis was performed using 6.0 N HCl (Sigma-Aldrich, USA, cat# 2104) at 110°C for 18 hours. Subsequently, neutralization was carried out using 5.7 N NaOH (VWR International, cat# BDH7247-1). If necessary, the samples were diluted with deionized water. 100 µL of the diluted samples and the standard OH-proline stock solutions were sequentially mixed with 0.05 N chloramine-T (Sigma-Aldrich, USA, cat# 857319-), 3.15 N perchloric acid (Sigma-Aldrich, USA, cat# 311421), and Ehrlich’s Reagent (Sigma-Aldrich, USA, cat# 03891). The absorbance of the resulting solution was measured at 560 nm. For the sGAG assay, dimethylmethylene blue (Sigma-Aldrich, USA, cat# 341088) solution was added to the diluted digested samples and chondroitin sulfate sodium salt (Sigma-Aldrich, USA, cat# C9819) solutions as standard reference. The sGAG content was determined by their absorbance at 525 nm, compared to those of reference solutions. For the DNA assay, 0.1 µg/mL Hoechst 33258 (Invitrogen, USA, cat# H3569) DNA stain in a buffer containing 10 mM Trizma base (Sigma-Aldrich, USA, cat# T6066), 0.1 mM NaCl, and 1 mM EDTA at pH 7.4 was added to the digested samples. The excitation and emission of samples were measured at 350 and 450 nm, respectively, and compared against a calf thymus DNA standard.

### Histological analysis

Post culture, hybrid tissues were washed with DPBS+/+, embedded in an optimal cutting temperature compound (Sakura, USA, cat# 4583), and flash-frozen in liquid nitrogen. Embedded samples were sectioned into 7 µm slices at -20°C, stained with Movat’s pentachrome (STTARR Innovation Centre, the University Health Network, Toronto), and scanned (Zeiss Axio Scan.Z1, Germany) by The Collaborative Advanced Microscopy Laboratories of Dentistry at the University of Toronto.

### Mechanical characterization

Hybrid structures were cut into 4.5×4.5 mm^2^ pieces using a parallel cutter, sprinkled with graphite powder for strain tracking, mounted to a biaxial mechanical tester (BioTester 5000, CellScale, Canada) using a tine sample attachment system (Biorakes with 0.7 mm tine spacing, CellScale, Canada) and stretched in equibiaxial loading cycles while submerged in a DPBS –/– bath at 37°C. Force and displacement values in both X and Y directions were collected, and the image tracking module of the BioTester software (LabJoy v.10.77) was used to track the X-Y coordinates of four points on the sample during the stretch cycles.

The original domain in the undeformed state of the sample was considered as − 𝐿⁄2 ≤ 𝑋𝑋_1_ ≤ 𝐿⁄2, − 𝐿⁄2 ≤ 𝑋𝑋_2_ ≤ 𝐿⁄2, and −𝐻⁄2 ≤ 𝑋𝑋_3_ ≤ 𝐻⁄2, where 𝐿 is the in-plane width and length of the square sample, H is its thickness, and 𝑋𝑋_1_ and 𝑋𝑋_2_ are equivalent to the X and Y coordinates, respectively. During the stretch cycles, the location of the tracked points were mapped to 𝑥_1_, 𝑥_2_, and 𝑥_3_ using 𝑥_1_ = 𝜆_1_𝑋𝑋_1_ + 𝐹_12_𝑋𝑋_2_, 𝑥_2_ = 𝐹_21_𝑋𝑋_1_ + 𝜆_2_𝑋𝑋_2_, and 𝑥_3_ = (ℎ⁄𝐻) 𝑋𝑋_3_, where 𝐹_12_ and 𝐹_21_ are the components of the transformation gradient tensor:

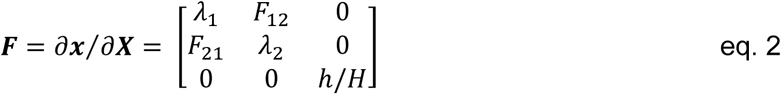

Assuming the test sample was incompressible, the determinant of the transformation gradient tensor was set to 1, and therefore, the thickness of the sample during deformation was calculated using ℎ = 𝐻⁄𝐽_2𝐷_, where 𝐽_2𝐷_ = 𝜆_1_𝜆_2_ − 𝐹_12_𝐹_21_. The method used to determine the components of 𝐹 using the coordinates of the tracked points on the test sample was described previously in detail by Labrosse *et al* ^23^. Knowing these components, the Green strain tensor was calculated using 𝐸 = 1⁄2 (𝐹^𝑇^. 𝐹 − 𝐼), where the dot represents the scalar product of two tensors. The components of the membrane tension tensor were calculated using 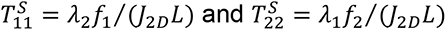 where 𝑓_1_ and 𝑓_2_ are the force readings in the X and Y directions, respectively. The resultant 𝑇^𝑆^ -𝐸_11_ and 𝑇^𝑆^ -𝐸_22_ curves were plotted to represent the tension-strain relationships in the X and Y directions of the hybrid tissue. The calculation of Green strain and membrane tension was performed using a MATLAB code.

### Parametric modelling

The relationship between polymeric scaffold architectural parameters and its biaxial mechanical properties was previously determined using a full factorial design of experiments and parametric modelling implemented in JMP Pro 17^25^. Knowing the mechanical properties of a polymeric scaffold and its corresponding hybrid tissue after three weeks of culture, the contribution of cell-laden hydrogel to the mechanical properties of the hybrid tissue was determined. A regression model was developed with scaffold fibre wave amplitude, 𝐴𝑚𝑝_𝑐𝑖𝑟𝑐._ and 𝐴𝑚𝑝_𝑟𝑎𝑑._, as factors and the deviation of hybrid tissue tension-strain behaviour from that of target native tissue, 𝑌_𝑐𝑖𝑟𝑐._ and 𝑌_𝑟𝑎𝑑._, as response variables according to the equations below.

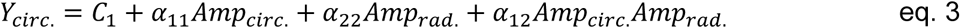

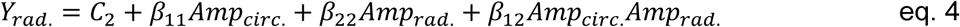

where 𝐶_𝑖_, 𝛼_𝑖𝑖𝑖_, and 𝛽_𝑖𝑖𝑖_ are the model parameters and 𝑌_𝑐𝑖𝑟𝑐._ and 𝑌_𝑟𝑎𝑑._ are defined based on the area between the tension-strain curve and the tension axis as shown below.

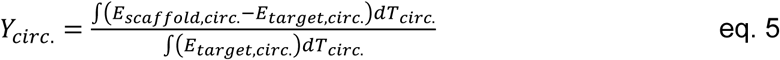

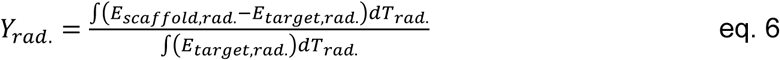

where 𝐸 and 𝑇 represent Green strain and membrane tension, respectively.

Model parameters, 𝛼_𝑖𝑖𝑖_ and 𝛽_𝑖𝑖𝑖_, were determined using least square estimation. When necessary, log transformation was performed to achieve normally distributed residuals, an essential assumption for least square regression. Via backward elimination, insignificant factors with a cut- off of p<0.1 were removed to obtain the reduced model, which was used to recalculate the parameters, 𝛼, and 𝛽. Finally, optimum wave amplitudes were determined using the Prediction Profiler in JMP with maximizing desirability, defined based on zero deviation of scaffold mechanics from that of native tissue. The deviation of the optimized scaffold tension-strain curve from that of associated native tissue was calculated using:

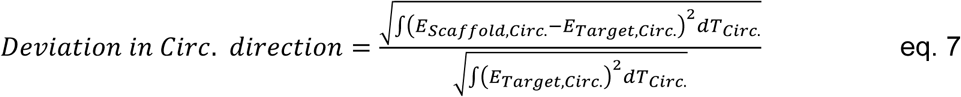

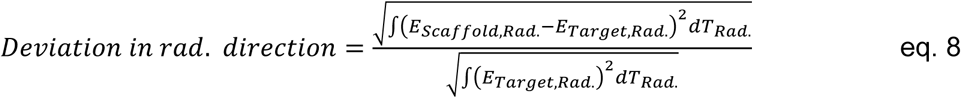

### Fabrication of trileaflet valves and hemodynamic testing

After three weeks of culture, hybrid tissues were wrapped around and sutured to a custom-designed crown-shaped frame, 3D printed using a biomedical-grade polymer (BioMed White, FormLabs, USA), to form a trileaflet valve with 12 mm of diameter. The trileaflet valve was mounted into a pulse duplicator bioreactor (Aptus Bioreactors, USA) and tested under controlled pulmonary arterial systolic pressure waveforms at 60 beat/min for 0.5×10^6^ cycles. Pressure and flow rate data were collected at every 300 cycles and processed using a MATLAB code to determine the average pressure profile as well as systolic and diastolic transvalvular pressures using the equations below.

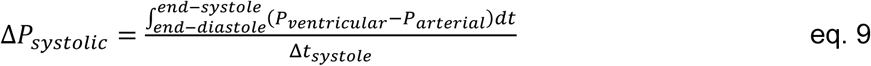

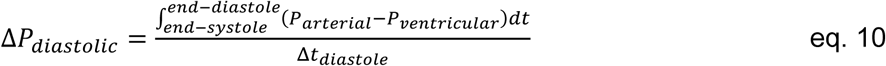

where 𝑃 is pressure, Δ𝑡_𝑠𝑠𝑠𝑠𝑡𝑜𝑙𝑒_ is the duration of systole, and Δ𝑡_𝑑𝑖𝑎𝑠𝑡𝑜𝑙𝑒_ is the duration of diastole.

### Diastolic test

A testing device was designed and developed via machining acrylic rods to expose the engineered PVs to diastolic transvalvular pressure in a static manner (**Figure S10A**). The outlets of the two sides of the device were connected to two reservoirs, placed at different heights, to generate physiological diastolic transvalvular pressure (10 mmHg), simulating the diastolic phase of the cardiac cycle, or at the same height to remove the pressure on the valve. The device was placed in the scanning chamber of a 𝜇CT system (Smart+ Irradiator, Precision, USA) and imaged (STTARR Innovation Centre, the University Health Network, Toronto) with 100-𝜇m resolution (**Figure S10B**). The resultant scan data were used to reconstruct the 3D model of the engineered PVs and determine leaflet belly curvature and strain by 3D Slicer and Fiji software. Leaflet belly curvature was calculated using the equation below.

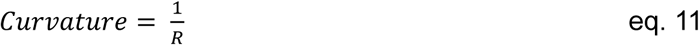

where 𝑅 is the radius of an arc coincident to the central belly region of the valve leaflet in its axial section.

Engineering strain in the radial and circumferential directions was calculated using the equation below.

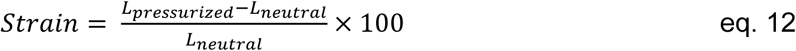

where engineering strain is calculated as percentage in the radial and circumferential direction, and 𝐿_𝑝𝑟𝑒𝑠𝑠𝑢𝑟𝑖𝑧𝑒𝑑_ and 𝐿_𝑛𝑒𝑢𝑡𝑟𝑎𝑙_ represent the radial or circumferential length of the leaflet at a section passing through the centre of the leaflet belly at 10 mmHg and 0 mmHg transvalvular pressure, respectively.

### Statistical analysis

Statistical analyses were performed using JMP Pro 17 software. Data are reported as mean ± standard deviation (SD). For pairwise comparison, data were analyzed using one-way ANOVA with Tukey’s post hoc test. Data were visualized using GraphPad Prism 9.

## Acknowledgements

This work was supported by the Canadian Institutes of Health Research [MOP-130481, CPG- 151962], the Natural Sciences and Engineering Research Council of Canada (NSERC) [CHRPJ 508364-17, RGPIN-2016-06026], the CONNECT! NSERC CREATE Program, the Translational Biology and Engineering Program of the Ted Rogers Centre for Heart Research, an NSERC Canada Graduate Scholarship – Doctoral and an Ontario Graduate Scholarship to BM, and an NSERC Postdoctoral Fellowship to NL.

