## Supplemental Information for "Biomimetic biaxial mechanical properties enhance hemodynamic performance and prevent adverse tissue remodelling in tissue-engineered heart valves"

**[#]Corresponding Authors**

### 1- Fabrication of hybrid tissue

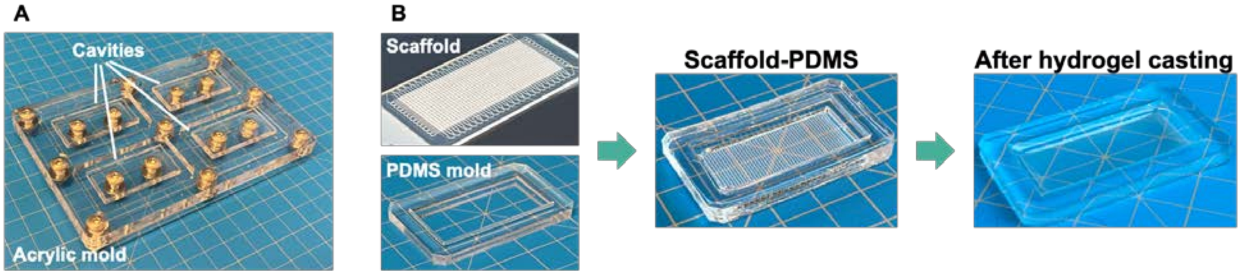


**Figure S1. A**) An acrylic mold to cast and fabricate polydimethylsiloxane (PDMS) structures. **B)** Polycaprolactone scaffolds were fabricated using melt electrowriting, attached to PDMS structures (molds), and seeded with human umbilical cord perivascular cells using fibrin hydrogel as a cell carrier.

### 2- Optimization of hydrogel parameters

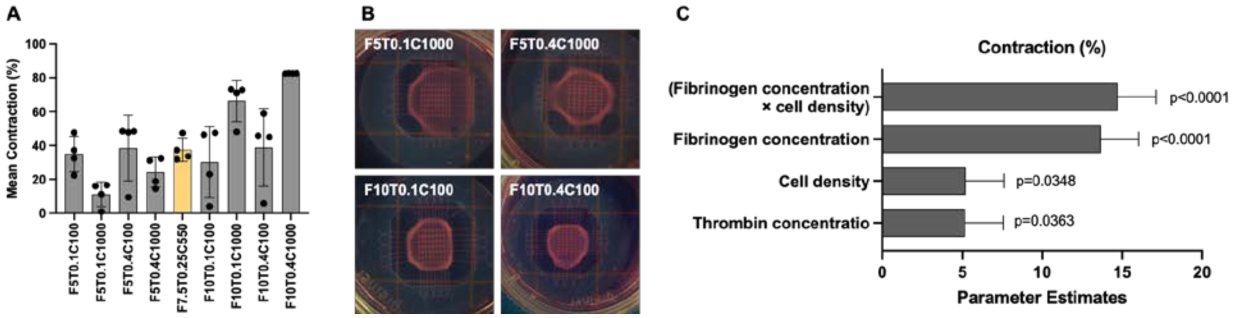


**Figure S2**. **A)** A full factorial design of experiments was performed to optimize fibrinogen (F) concentration, thrombin (T) concentration, and cell (C) density in the cell-lade fibrin hydrogel to minimize their contraction during the static culture. Fibrinogen concentration varied from 5 mg/ml (F5) to 10 mg/ml (F10); thrombin concentration varied from 0.1 unit/ml (T0.1) to 0.4 unit/ml (T0.4); cell density varied from $100\times10^3$ cell/ml (C100) to $1000\times10^3$ cell/ml (C1000). **B)** Microscopic images of hybrid tissues with various compositions after three weeks of culture. **C)** Parameter estimate calculated using least square regression to relate tissue contraction and hydrogel composition. These estimations were used to determine the composition of cell-laden fibrin hydrogel that yielded minimum contraction during three weeks of culture.

#### 3- Determination of engineered tissue maturation time required for valvular function

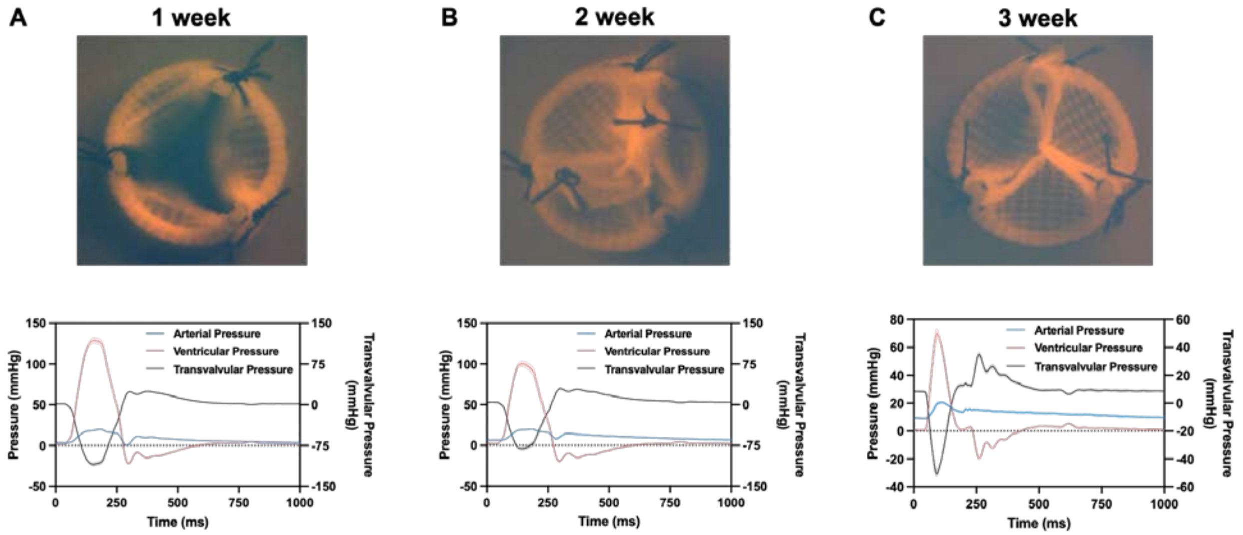


**Figure S3.** Engineered pulmonary valves, built from hybrid tissues matured for one (**A**), two (**B**), or three (**C**) weeks, under physiological pulmonary conditions and their associated pressure profiles for a single cardiac cycle. Images were taken at day two for (A) and (B) and at day five for (C).

### *4-* Modification to the engineered valve supporting crown to increase cell viability

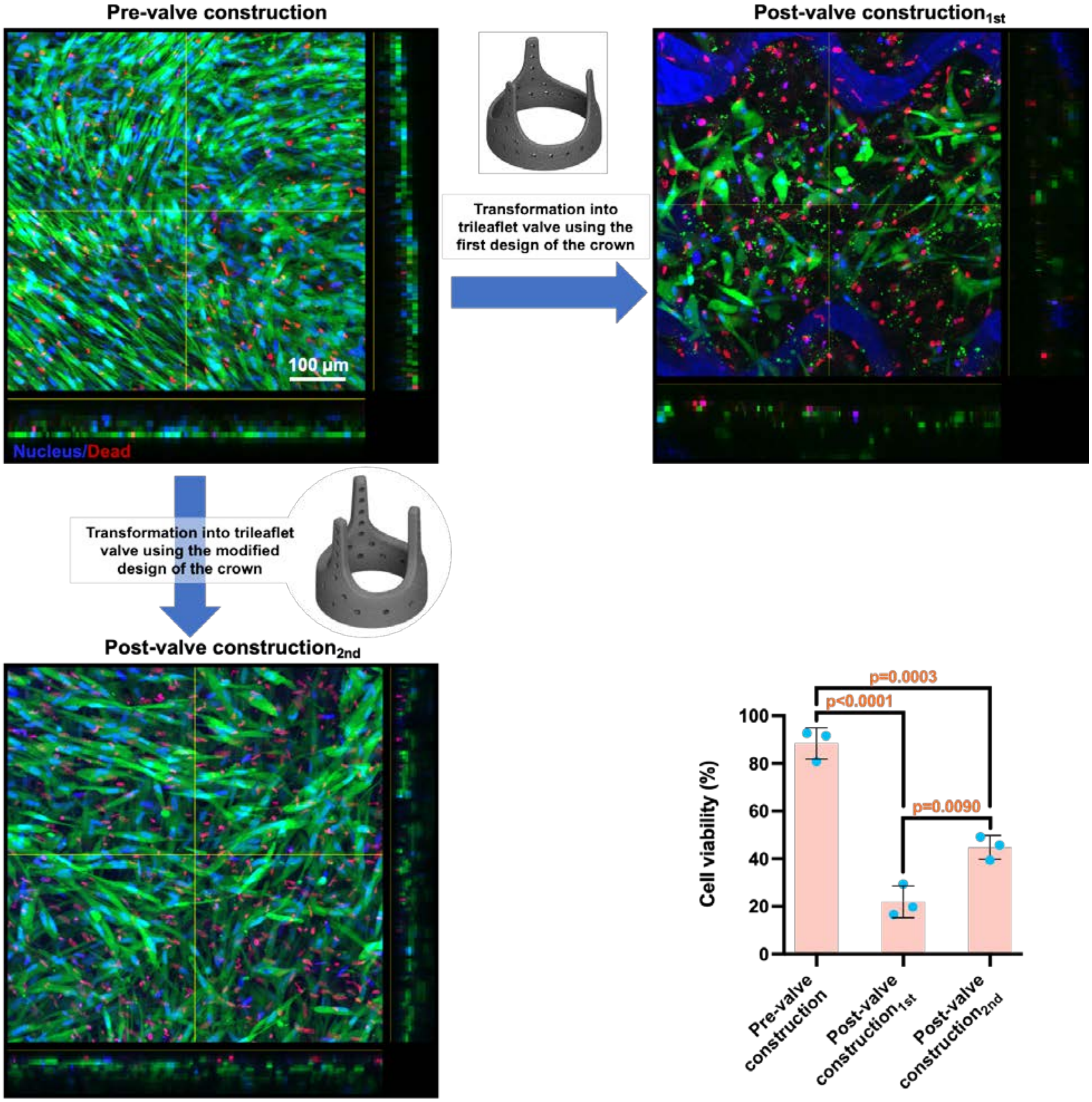


**Figure S4.** The viability of hybrid tissue after trileaflet valve assembly using two different designs of the crown-shaped frame. The second design has fewer suture sites with a larger diameter.

5- **Analysis of dead cell nuclei size in engineered hybrid tissues at various stages: before and after transformation into trileaflet valve, and after dynamic culture**

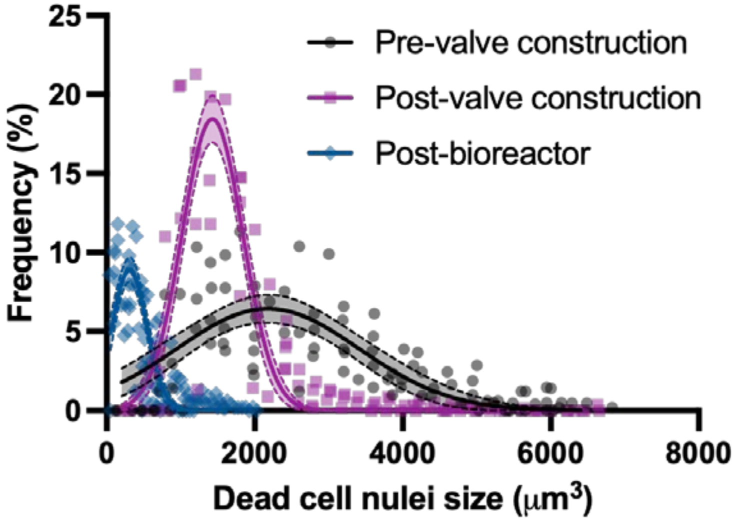


**Figure S5.** Size distribution of dead cell nuclei in hybrid tissues pre- and post-trileaflet valve construction and post-bioreactor, after valvular function for $0.5 \times 10^6$ cycles.

### 6- Gross structure of the polychrome scaffold constituent of the engineered pulmonary valves (PVs) after dynamic culture

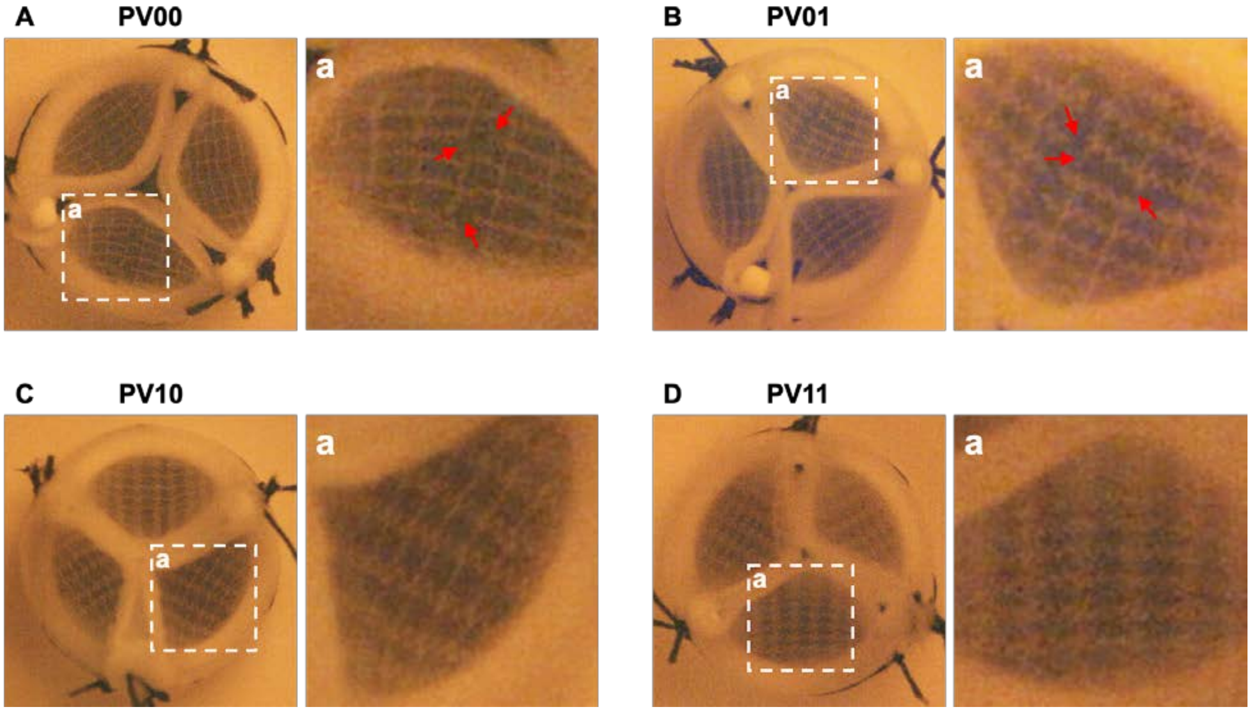


**Figure S6.** Engineered valves (PVs) under the physiological pulmonary pressure and flow rate and the condition of their constituent polymeric scaffold. **A)** PV00 with non-native mechanics in both radial and circumferential directions showed some damage to the polymeric scaffold after valvular function. **B)** PV01 with native mechanics in circumferential but non-native mechanics in radial direction showed some damage to the polymeric scaffold after valvular function. **C)** PV10 with native mechanics in radial but non-native mechanics in circumferential direction showed intact fibre morphology after valvular function. **D)** PV11 with native mechanical properties in both radial and circumferential directions showed intact fibre morphology after valvular function.

### 7- Determination of the crystallinity of polycaprolactone (PCL) constituent of engineered PVs

Samples of engineered PV leaflets from all four valve types, before and after dynamic culture, were digested via incubating in 1 mg/ml collagenase type I (Sigma-Aldrich, USA, cat# C0130) solution in Dulbecco's phosphate-buffered saline (DPBS)−/− (Gibco, USA, cat# 14190-144) at 37 °C for 5 hours and, subsequently, at room temperature overnight. The digestion process removed the tissue part, leaving the melt electro-written PCL scaffolds for determination of polymer crystallinity using a Mettler-Toledo DSC1 differential scanning calorimeter. The samples were run at a heating and cooling rate of 10 °C/min that was applied over a temperature range program containing two cycles between −100 and 100 °C. $\Delta H_f$ was determined by calculating the area under the melting endotherm and dividing it by the mass of the test sample. The degree of crystallinity, $\%X$, was determined using the equation below:

$$\%X = \frac{\Delta H_f}{\Delta H_f^{\circ}}$$

where $\Delta H_f^{\circ}$ is the enthalpy of fusion for 100% crystalline PCL, which was considered 139.5 mJ/mg as determined previously[1].

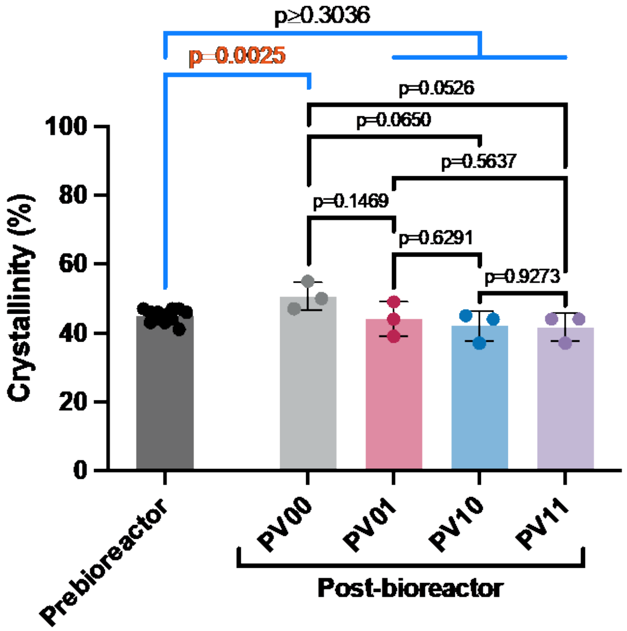


**Figure S7.** The crystallinity of polycaprolactone in the polymeric scaffold constituent of the four types of pulmonary valves (PVs) before and after 5 days of valvular function.

### *8-* Bending behaviour of the hybrid tissues

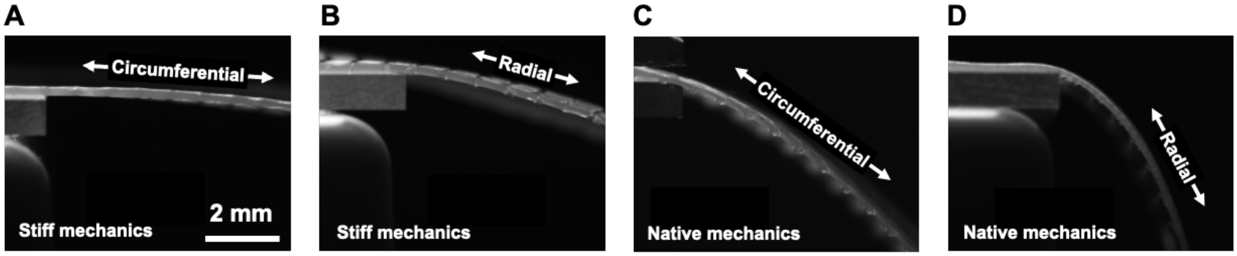


**Figure S8.** Microscopic images of three-week matured hybrid tissues with stiff circumferential (**A**), stiff radial (**B**), native circumferential (**C**), or native radial (**D**) mechanical properties, showing their bending properties while an equal length of each tissue (10 mm) was hanging over the free edge of a glass slide.

### 9- Diastolic and systolic function of the engineered PVs compared to healthy and disease physiological levels

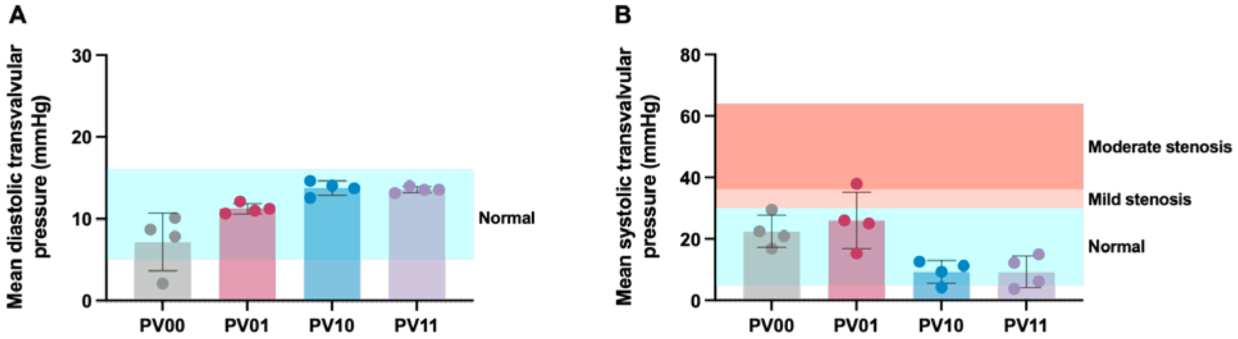


**Figure S9**. Diastolic (**A**) and systolic (**B**) transvalvular pressure of the four types of engineered pulmonary valves compared to native healthy and diseased levels.

### *10-*Design of the testing device developed to enable computed tomography scanning of engineered PVs while under physiological diastolic pressure

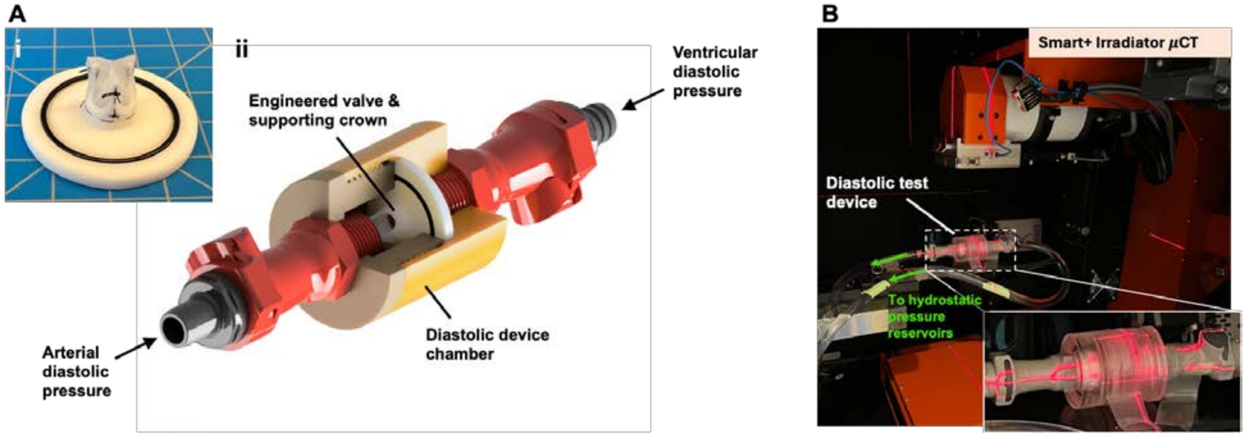


**Figure S10. A)** Trileaflet pulmonary valve constructed using a three-week matured hybrid tissue and a supporting crown for diastolic pressure test (i). Schematic of the developed diastolic test device to expose the engineered pulmonary valve to the physiological diastolic transvalvular pressure in a static manner (ii). **B)** The diastolic test device in a micro-computed tomography ($\mu$CT) device to image the engineered pulmonary valve under the physiological diastolic transvalvular pressure.
